# Identification and Characterization of ML321: a Novel and Highly Selective D*2* Dopamine Receptor Antagonist with Efficacy in Animal Models that Predict Atypical Antipsychotic Activity

**DOI:** 10.1101/2022.11.14.516475

**Authors:** R. Benjamin Free, Ashley N. Nilson, Noelia M. Boldizsar, Trevor B. Doyle, Ramona M. Rodriguiz, Vladimir M. Pogorelov, Mayako Machino, Kuo Hao Lee, Jeremiah W. Bertz, Jinbin Xu, Herman D. Lim, Andrés E. Dulcey, Robert H. Mach, James H. Woods, J Robert Lane, Lei Shi, Juan J. Marugan, William C. Wetsel, David R. Sibley

## Abstract

We have developed and characterized a novel D2R antagonist with exceptional GPCR selectivity – ML321. In functional profiling screens of 168 different GPCRs, ML321 showed little activity beyond potent inhibition of the D2R, and to a lesser extent the D3R, demonstrating excellent receptor selectivity. The D2R selectivity of ML321 may be related to the fact that, unlike other monoaminergic ligands, ML321 lacks a positively charged amine group and adopts a unique binding pose within the orthosteric binding site of the D2R. PET imaging studies in non-human primates demonstrated that ML321 penetrates the CNS and occupies the D2R in a dose-dependent manner. Behavioral paradigms in rats demonstrate that ML321 can selectively antagonize a D2R-mediated response (hypothermia) while not affecting a D3R-mediated response (yawning) using the same dose of drug, thus indicating exceptional *in vivo* selectivity. We also investigated the effects of ML321 in animal models that are predictive of antipsychotic efficacy in humans. We found that ML321 attenuates both amphetamine- and phencyclidine-induced locomotor activity and restored pre-pulse inhibition (PPI) of acoustic startle in a dose-dependent manner. Surprisingly, using doses that were maximally effective in both the locomotor and PPI studies, ML321 was relatively ineffective in promoting catalepsy. Kinetic studies revealed that ML321 exhibits slow-on and fast-off receptor binding rates, similar to those observed with atypical antipsychotics with reduced extrapyramidal side effects. Taken together, these observations suggest that ML321, or a derivative thereof, may exhibit “atypical” antipsychotic activity in humans with significantly fewer side effects than observed with currently FDA-approved D2R antagonists.

---

G protein–coupled receptors (GPCRs) are the largest family of cellular receptors in mammals and are important targets for approximately one-third of all FDA–approved drugs^1^. These receptors regulate multiple physiological processes by transducing extracellular stimuli, such as neurotransmitters, hormones, peptides, or light, into intracellular signals through activating both G protein–dependent and G protein–independent pathways, leading to second messenger generation and downstream signaling events^2^. Receptors for the neurotransmitter dopamine constitute a subfamily of GPCRs that are critically important for the regulation of movement, learning, mood, reward, and attention within the central nervous system. Dopamine receptors are divided into two subfamilies, D1-like (D1R and D5R) and D2-like (D2R, D3R, and D4R) on the basis of their sequence homology, signaling pathways, and pharmacological properties^3–5^. Amongst these receptors, the D2R subtype is an extremely well validated drug target for the therapy of Parkinson’s disease, schizophrenia, and other neuropsychiatric disorders^6^. Surprisingly, there are relatively few drugs with high selectivity for the D2R despite efforts from focused chemistry campaigns and the obvious clinical need for such agents. This arguably relates to the high structural conservation of the orthosteric binding site of the D2R with that of other dopamine receptors and related GPCRs^7, 8^.

Schizophrenia is a devastating illness that affects approximately 1.1% of the adult human population^9^. It is characterized by a combination of positive symptoms (hallucinations, delusions), negative symptoms (social withdrawal), and cognitive deficits. The drugs currently used to treat schizophrenia are classified as either typical (first-generation) or atypical (second-generation) antipsychotics^10^, which share the feature of D2R antagonism. This nomenclature stems from the fewer extrapyramidal symptoms (EPS), or parkinsonian-like side effects, observed with atypical antipsychotics. So-called “third generation” antipsychotics have recently been developed that are low-efficacy partial D2R agonists with atypical side-effect profiles, although these exhibit an increased risk of akathisia, potentially due to D2R stimulation^11^. While other drug targets are being evaluated for the treatment of schizophrenia^12–15^, particularly for ameliorating negative and cognitive symptoms, D2R antagonism remains the primary mechanism underlying current antipsychotic medications. Notably, however, pimavanserin, an inverse agonist at the 5-HT_2A_ serotonin receptor, has been approved for treating psychosis associated with Parkinson’s disease^16^ and may be effective as an adjunctive therapeutic (combined with an antipsychotic) for the treatment of negative symptoms in schizophrenia^17^.

Despite considerable research, the mechanism of “atypicality”, or decreased EPS, observed with second or third generation antipsychotics is unknown. An older hypothesis suggested that this may be due to 5-HT_2A_ serotonin receptor blockade^18^. More recent data, however, suggest that receptor binding kinetics and duration of D2R occupancy may be more relevant^19, 20^. There is consensus, however, that attenuation of D2R signaling is responsible for the efficacy of these drugs in treating the core (positive) symptoms of schizophrenia^10, 21–24^. Unfortunately, all antipsychotics, both typical and atypical possess numerous other side effects (sedation, weight gain, diabetes, etc.), primarily due to their off-target interactions with other receptors and drug targets^25–27^. Thus, a globally selective D2R antagonist, which has not been previously available, could be particularly effective for treating schizophrenia while exhibiting fewer off-target side effects.

Recently, we conducted a high throughput screen of the NIH small molecule repository and identified a small pool of compounds that were more than 10-fold selective for the D2R *versus* the D3R. As the D3R is the receptor most closely related to the D2R, both structurally and pharmacologically, we thought it would be advantageous to chemically optimize one of the D2R-selective hit compounds to improve both its potency and selectivity. Using D2R>D3R selectivity to drive the chemistry, we identified a lead chemical probe compound, ML321, and described its corresponding structure-activity relationships (SAR)^28^. ML321 was shown to exhibit >40-fold selectivity for the D2R *versus* the D3R using radioligand binding and functional assays^28^. In the current study, we show that ML321 has unprecedented selectivity for the D2R compared to many other biogenic-amine and related GPCRs. Using molecular modeling and mutagenesis approaches, we find that ML321 adopts a unique binding pose within the orthosteric binding site of the D2R and we further demonstrate that it functions as an inverse agonist at this receptor. Behavioral paradigms in rodents demonstrate that ML321 can selectively antagonize a D2R-mediated response while not affecting a D3R-mediated response; thereby indicating excellent *in vivo* selectivity. ML321 was further shown to be active in animal models that are predictive of antipsychotic efficacy in humans. Importantly, it was found to be relatively ineffective in producing catalepsy in rodents, which may be explained by the fact that ML321 exhibits slow-on and fast-off receptor binding rates, similar to those of atypical antipsychotics. The exceptional receptor selectivity and unique binding kinetics of ML321 suggest that it will serve as a useful chemical probe for the D2R and as an advanced drug lead for an improved antipsychotic with decreased on- and off-target side effects when compared to existing antipsychotic drugs in current in use.

## RESULTS AND DISCUSSION

### ML321 Exhibits Unprecedented Selectivity as a D2R Antagonist

As discussed, the existing pharmaceutical armamentarium is lacking in highly selective D2R antagonists, especially those that can differentiate the D2R from the D3R^6^. Thus, to identify novel and more selective D2R antagonist scaffolds, we initiated a high throughput screening campaign which initially identified a small pool of hit compounds that were >10-fold D2R/D3R selective^28^. The most promising antagonist candidate was further optimized using medicinal chemistry to increase the D2R *versus* D3R selectivity resulting in the lead chemical probe compound ML321^28^. The structure of ML321 is shown in **Figure 1A**. One notable feature of ML321 is that it lacks a protonatable nitrogen (at physiological pH) found in almost all biogenic amine receptor ligands, which typically forms an ionic bond with a conserved aspartate residue (Asp^3.32^, using the numbering system of Ballesteros and Weinstein^29^) within the receptor’s orthosteric binding site^30^. Another unique feature is the chiral sulfoxide moiety in the benzothiazepine ring. ML321 consists solely of the *S*-enantiomer of this sulfoxide as we have previously determined that the *R*-enantiomer lacks affinity for the D2R^28^. To examine ML321 selectivity among all dopamine receptor subtypes, we initially performed radioligand binding competition assays (**Figure 1B).** We found that ML321 has the highest affinity for the D2R with a K_i_ value of 58 nM followed by the D3R with an approximated K_i_ value of 4 μM resulting in a D2R/D3R selectivity of ∼80-fold. In contrast, the D1R, D4R, and D5R all possess negligible affinities (K_i_ values >10 μM) for ML321.

**Figure 1.**
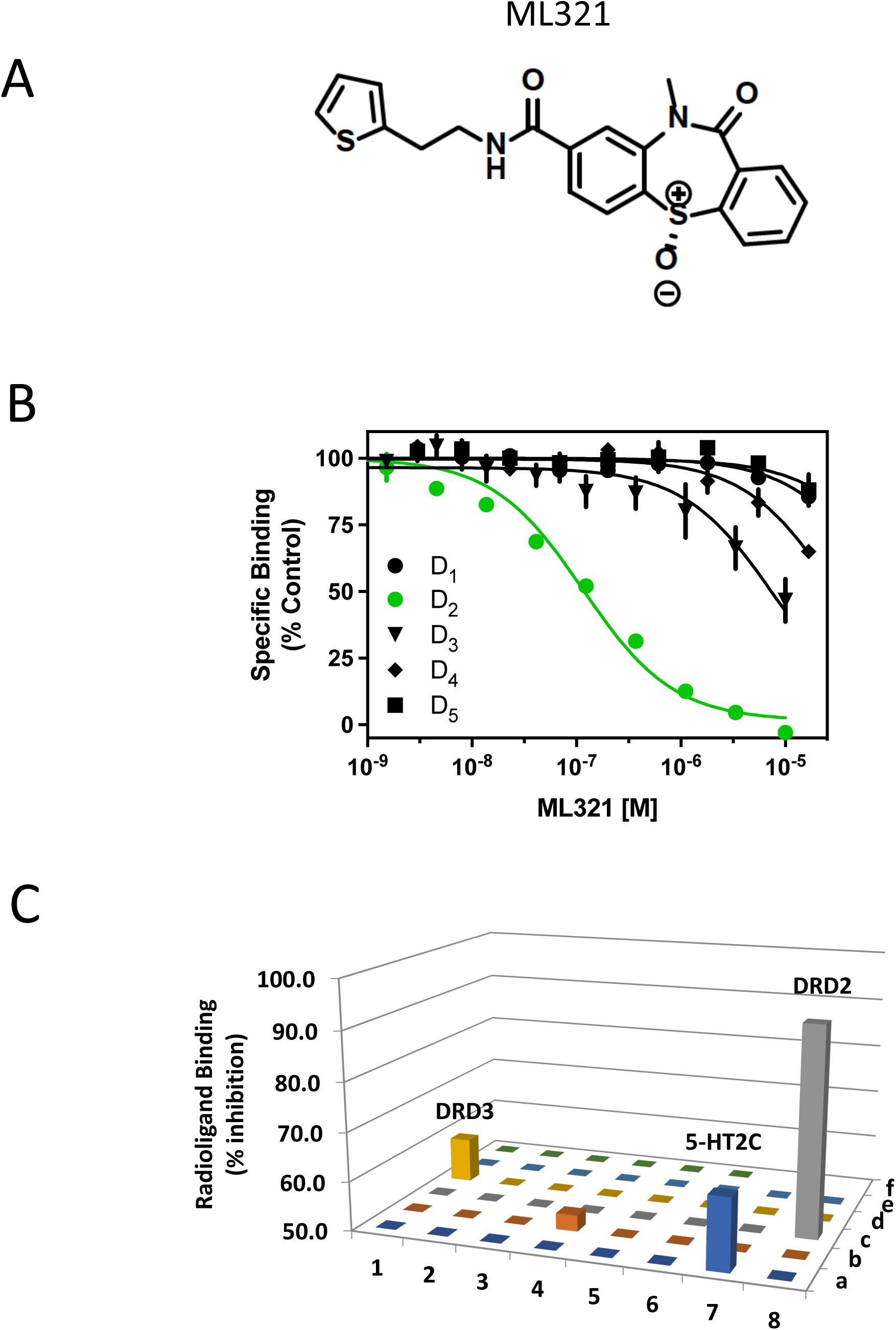
ML321 shows high selectivity for the D2R among dopamine and related biogenic amine receptors. (**A**) Structure of ML321. (**B**) Radioligand binding competition assays to determine ML321 affinity for dopamine receptors. Radioligand binding assays were performed as described in the Methods. The data are expressed as percentage of the control specific binding and represent mean ± SEM values from three independent experiments each performed in triplicate. Mean ML321 Ki values [95% C.I.] for each receptor were calculated from the IC_50_ values using the Cheng-Prusoff equation^31^ and found to be 57.6 nM [45.7-72.8] for the D2R and 3.9 μM [2.8-5-5] for the D3R. (Because the curve for the D3R was not complete, the Ki for ML321 should be considered an approximation). The IC_50_ values for the D1R, D4R and D5R were >10 μM. (**C**) Psychoactive Drug Screening Program (PDSP) results. Radioligand binding assays were performed as described in the Methods. Each bar represents a unique drug target (**Table S1**). This process identified four receptors for which 10 µM of ML321 inhibited the specific binding by more than 50% (defined as the level of significance by the PDSP): D2R (92%) and D3R (59%), 5-HT_2C_ (64%) and 5-HT_7_ (53%) serotonin receptors.

To further assess ML321 selectivity amongst a larger group of drug targets, we used the NIMH Psychoactive Drug Screening Program (PDSP), which utilizes radioligand binding competition assays for 46 unique GPCRs, transporters and ion channels^32^. A complete list of these targets is listed in **Table S1**. For screening, a single high concentration (10 µM) of ML321 was used to compete for radioligand binding to the PDSP targets (**Figure 1C**). This process identified four receptors for which 10 µM of ML321 inhibited the specific binding by more than 50% (defined as the level of significance by the PDSP), which were (with % inhibition): the D2R (92%), D3R (59%), 5-HT_2C_ (64%) and 5-HT_7_ (53%) serotonin receptors. Since the screening concentration (10 µM) of ML321 in this assay was approximately 100-fold greater than the concentration needed to engage the D2R for clinical therapeutics (∼100 nM, 60-75% occupancy) ^33^, it is unlikely that the D3R or 5-HT_2C_ and 5-HT_7_ serotonin receptors would be occupied to a significant extent in clinical practice.

As the PDSP screening panel relies solely on radioligand binding to assess receptor selectivity, we utilized the DiscoverX gpcrMAX^TM^ functional assay, which measures agonist-stimulated β-arrestin recruitment to 168 known GPCRs (http://www.DiscoverX.com). As with the PDSP, we performed this screen using a high concentration (10 μM) of ML321 to maximize the detection of off-target activities. **Figure 2A** shows the results of this screen conducted in antagonist mode where each GPCR is stimulated with an EC_80_ concentration of a reference agonist, plus 10 μM ML321, followed by measurement of β-arrestin recruitment. ML321 maximally inhibited the short (D2R_S_) and long (D2R_L_) isoforms of the D2R, while partial inhibition (72%) of the D3R was observed at this high ML321 concentration. All other GPCRs were inhibited by less than 50%, although a few showed low (>20%), but significant inhibition including the 5-HT_2A_ and 5-HT_2C_ serotonin, BLT1 (leukotriene B_4_), sphingosine-1-phosphate 4, and α_2C_-adrenergic receptors. We also conducted the screen in agonist mode where each GPCR was treated with 10 μM ML321 (**Figure 2B**). As can be seen, the only GPCR that exhibited a response was the CB2 cannabinoid receptor (44% stimulation). Numerical results for **Figure 2** are provided in **Table S2**, whereas the data are also displayed in a heat-map format in **Figure S1.** As noted above, other than the D2R, it is unlikely that any of the other receptors tested here would be occupied to a significant extent when using therapeutically-relevant concentrations (100-250 nM) of ML321 at would occupy the D2R by 60-75%, which is the target occupancy for treating schizophrenia^33^. Taken together, these results demonstrate that ML321 is an exceptionally selective D2R antagonist.

**Figure 2.**
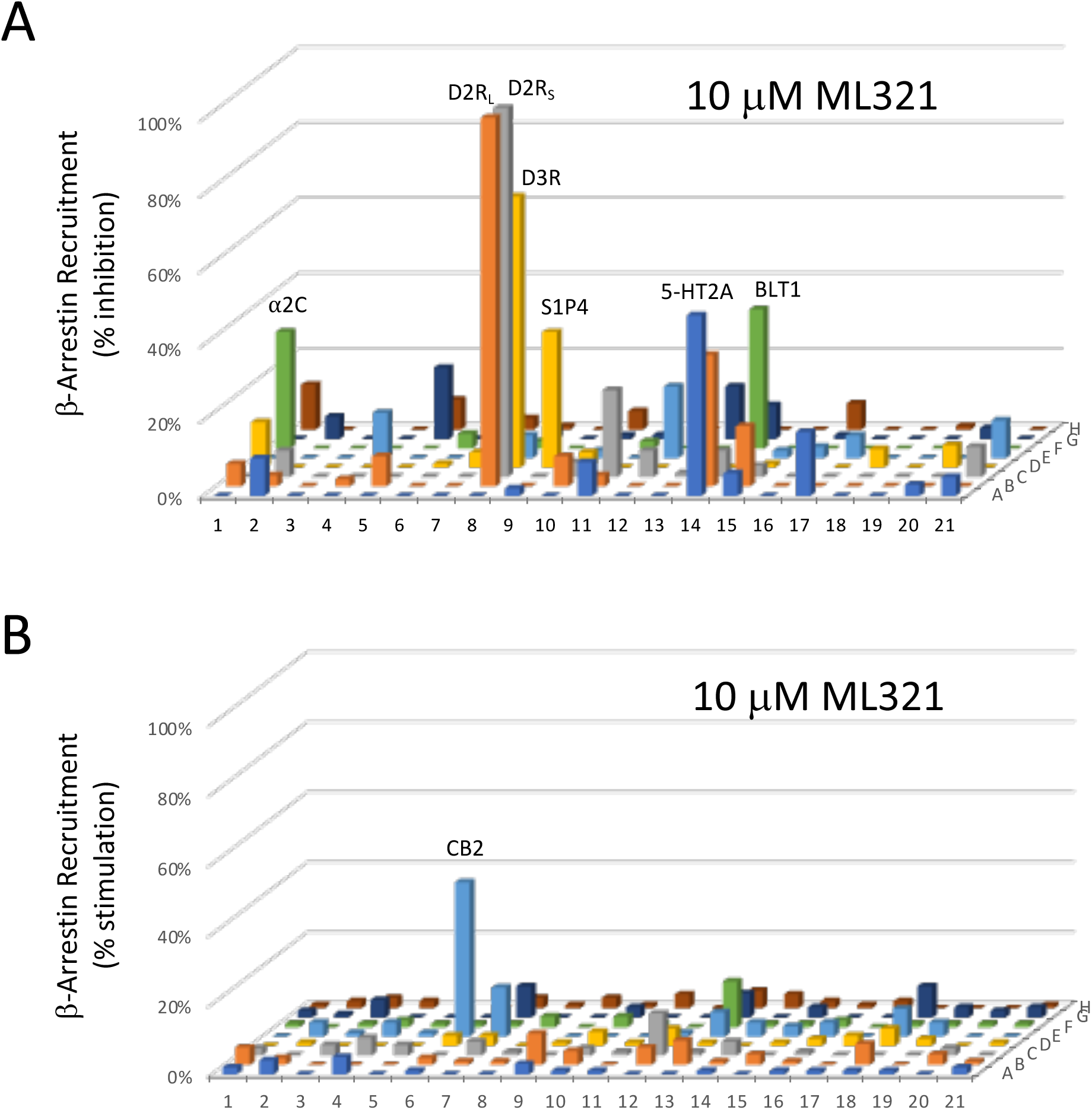
Functional profiling of ML321 against an array of 168 known GPCRs. A single high concentration (10 µM) of ML321 was screened using a β-arrestin recruitment assay in the DiscoverX gpcrMAX^TM^ assay panel in both antagonist (**A**) or agonist (**B**) modes as described in the Methods. Data represent the percent maximum stimulation (agonist mode) observed by a reference agonist for each GPCR or the percent inhibition (antagonist mode) of a response produced by an EC_80_ concentration of a reference agonist for each GPCR. A complete key to the GPCR array and numerical results are provided in **Table S2**. Only responses that deviated >20% from baseline are considered to be significant. In panel **A**, these include the following receptors (% activity): D2R_L_ (98%), D2R_S_ (98%), D3R (72%), 5-HT_2A_ (48%), 5-HT_2C_ (35%), BLT1 (leukotriene B_4_) (LTB4R) (37%), sphingosine-1-phosphate 4 (S1PR4) (36%), α_2C_-adrenergic (31%); and in panel **B**, CB2 cannabinoid (44%).

### ML321 Antagonizes the D2R in a Competitive Manner and Exhibits Inverse Agonist Activity

We next wished to characterize the nature of ML321’s antagonism of the D2R. We initially performed curve-shift experiments in which the ability of increasing concentrations of ML321 to modulate dopamine potency and efficacy were measured using different functional outputs. **Figure 3A** shows concentration-response curves (CRCs) for dopamine stimulation of β-arrestin recruitment in the presence of increasing concentrations of ML321. As can be seen, increasing concentrations of ML321 result in a parallel shift to the right of the dopamine CRCs such that there is an apparent decrease in dopamine potency (EC_50_) with no change in efficacy (E_max_). These data were fit to a Gaddum/Schild model of competitive antagonism resulting in a Schild slope of unity (see inset) allowing us to derive the affinity (*K*_B_) of ML321 for inhibiting this response, which was 100 nM – similar to the Ki for ML321 observed in the radioligand binding assays (**Figure 1**). We performed similar experiments using D2R modulation of forskolin-stimulated cAMP accumulation, which is G protein-mediated, as shown in **Figure 3B**. Using this functional output, increasing concentrations of ML321 also promote a parallel shift to the right in the dopamine CRCs consistent with a competitive form of antagonism.

**Figure 3.**
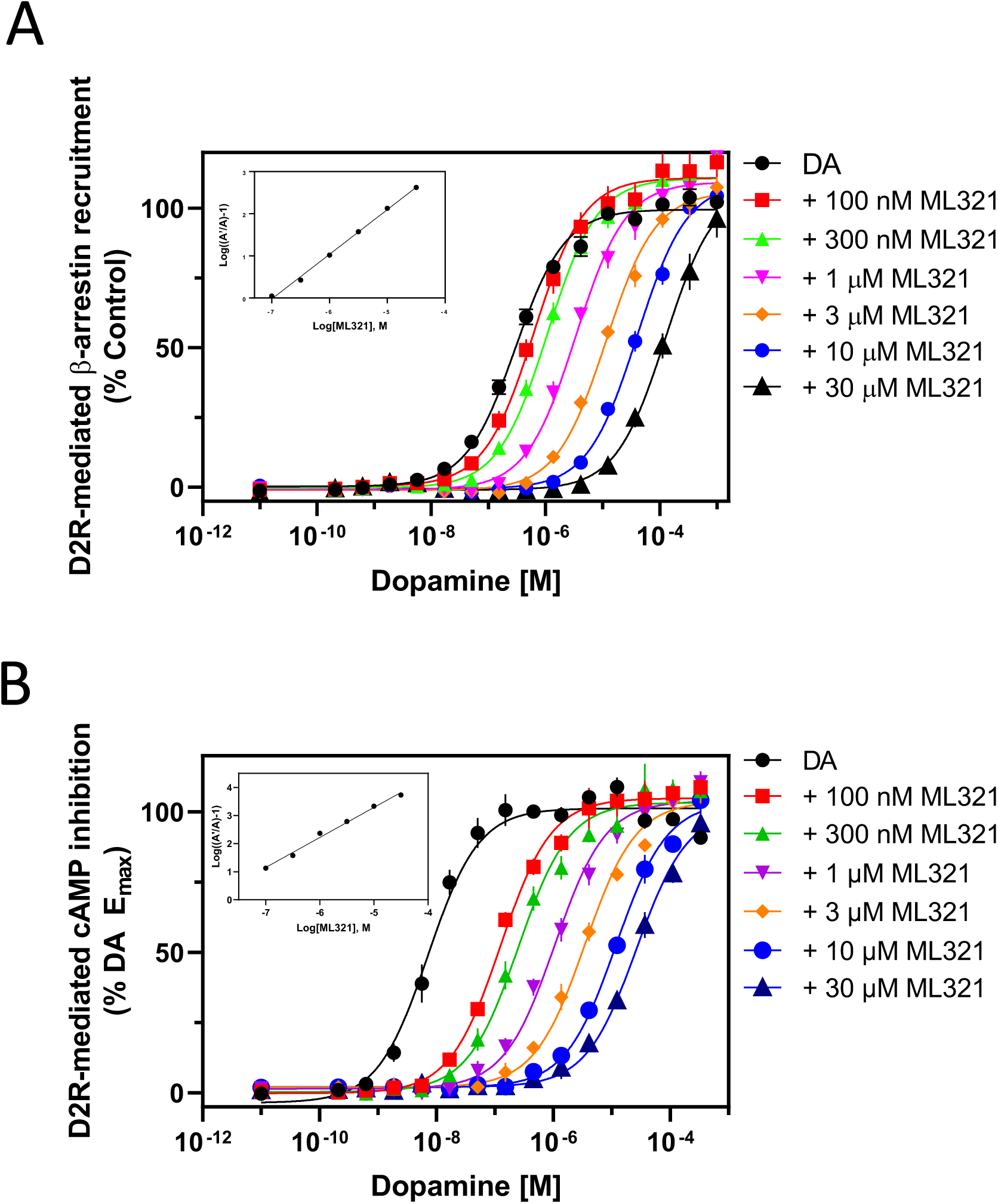
Curve-shift assays indicate that ML321 behaves in a competitive manner with dopamine at the D2R. D2R-mediated β-arrestin recruitment assays (**A**) or cAMP inhibition assays (LANCE) (**B**) were conducted by stimulating the receptor with the indicated concentrations of dopamine with or without various concentrations of ML321 as described in the Methods. For the cAMP assay, the cells were incubated with 10 μM forskolin to stimulate cAMP production. Data are expressed as a percentage of the maximum dopamine response seen in the absence of ML321 (% control) and represent the mean ± SEM of at least three independent experiments each performed in triplicate. Insets show Schild analyses of the data from which mean *K*_B_ values [95% C.I.] were derived. (**A**) ML321 *K*_B_ = 103 nM [76.7-127] (n=3), (**B**) ML321 *K*_B_ = 8.36 nM [3.5-16.5] (n=3).

Gaddum/Schild analyses of these data resulted in a *K*_B_ of 8.35 nM, which was more potent than that observed in the β-arrestin recruitment assay. The reason for this difference is not immediately obvious but could be due to the complexity of the cAMP assay, which involved stimulation of cAMP accumulation by forskolin, inhibition of this response by dopamine, and then reversal of the dopamine response by ML321.

We used two additional functional assays to assess ML321 antagonism of dopamine signaling through the D2R. The first was a BRET-based D2R-mediated β-arrestin recruitment assay shown in **Figure S2A**. In this assay, a BRET signal is produced when dopamine stimulates β-arrestin recruitment to the D2R. Here, too, we found that increasing ML321 concentrations in the assay promoted an apparent decrease in dopamine potency with no change in efficacy for promoting β-arrestin recruitment. The mean *K*_B_ for ML321 in this assay was 230 nM. Finally, we used a BRET-based Go activation assay, as shown in **Figure S2B**, where the BRET donor and acceptor pairs are fused to the α and γ subunits of the Go heterotrimer, respectively, such that a constitutive BRET signal is produced. D2R-mediated activation of Go promotes dissociation of the α subunit from the βγ subunits resulting in a decrease in BRET signal. Despite the relatively small BRET signal, we were able to establish that, as with the other functional assays, that ML321 behaves as a competitive antagonist with a mean *K*_B_ of 143 nM. While the observed potencies of ML321 were somewhat variable across the four functional assays examined, all of them served to establish that ML321 is a competitive antagonist at the D2R.

The D2R has been shown to exhibit constitutive activity in various functional signaling outputs^34^ and many antagonists of the D2R exhibit inverse agonist activity to varying degrees^35^. Thus, we decided to investigate the possibility that ML321 displays inverse agonist activity at the D2R using the Go BRET activation assay as shown in **Figure 4**. As described above, in the presence of the D2R, the modified Go heterotrimer exhibits constitutive BRET activity in the absence of agonist, which is defined as the baseline (zero) signal in this assay. Dopamine stimulation of the D2R promotes Go activation (heterotrimer dissociation) and a decrease in the BRET signal (**Figure 4**). In contrast, saturating concentrations of various D2R antagonists including sulpiride, eticlopride, spiperone, or butaclamol increased the BRET signal through inverse agonist activity at the D2R and stabilization of the Go heterotrimer. ML321 also increased the BRET signal, suggesting that it exhibits inverse agonist activity to an extent that was indistinguishable from that of the other antagonists (**Figure 4**). Moreover, we established that the inverse agonist activity of ML321 was dose-dependent, as illustrated in **Figure S3.** In this experiment, ML321 exhibited an EC_50_ of 275 nM for the response, although the data were quite variable given the extremely small signal observed in this BRET output.

**Figure 4.**
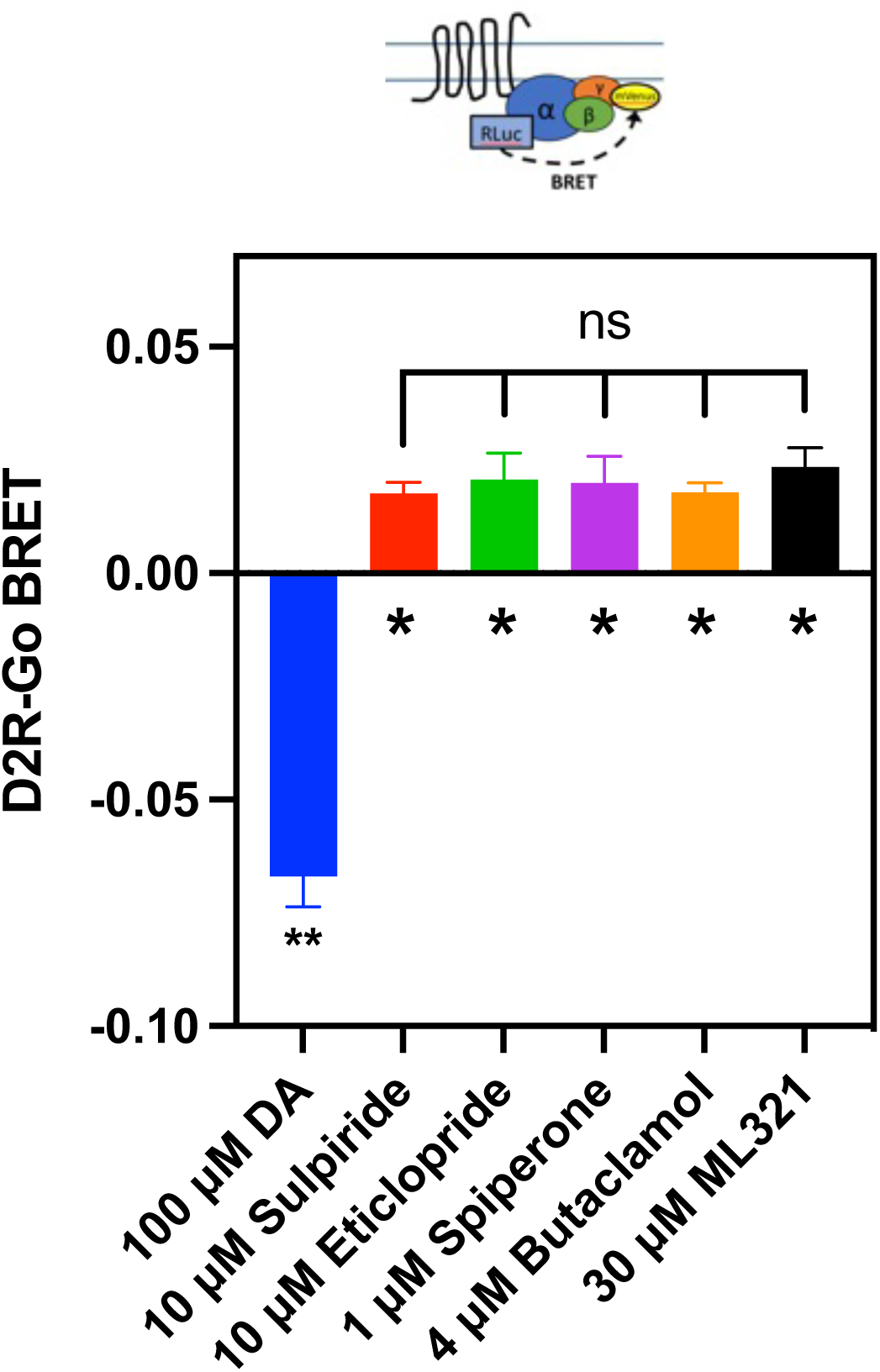
ML321 exhibits inverse agonist activity at the D2R. Go BRET activation assays were performed as described in the Methods. Data represent the mean ± SEM of 4-6 independent experiments each performed in octuplicate and are expressed as Δ BRET from baseline, which represents the constitutive BRET signal seen in the absence of any drug treatment. Dopamine produces a dissociation of the Go heterotrimer and thus a decrease in BRET, whereas due to constitutive activity of the D2R, the tested antagonists exhibit inverse agonist activity through the promotion of heterotrimer formation. The Δ BRET signals for each compound were statistically different from baseline as determined using Dunnett’s multiple comparisons test following a one-way ANOVA (* indicates p < 0.05 and ** indicates p < 0.0001). The Δ BRET signals for the antagonists did not statistically differ from each other (p = 0.841 using one-way ANOVA).

### ML321 Adopts a Unique Binding Pose Within the Orthosteric Site of the D2R

To characterize the binding mode of ML321 at the D2R, we performed a computational modelling and simulation study to investigate the interactions between ML321 and the D2R. Based on the previous SAR study of the ML321 scaffold^28^, we hypothesized that the dibenzothiazepine moiety of ML321 orients within the orthosteric binding site (OBS) of the D2R, while the thiophene moiety is oriented towards a secondary binding pocket (SBP). From the docking results in our D2R model, we selected several binding poses of ML321 consistent with this hypothesis and conducted molecular dynamics (MD) simulations (see Methods) for each of the resulting complexes. During these simulations, the various docking poses converged onto a single binding pose within the OBS (**Figure 5A**), which is described below.

**Figure 5.**
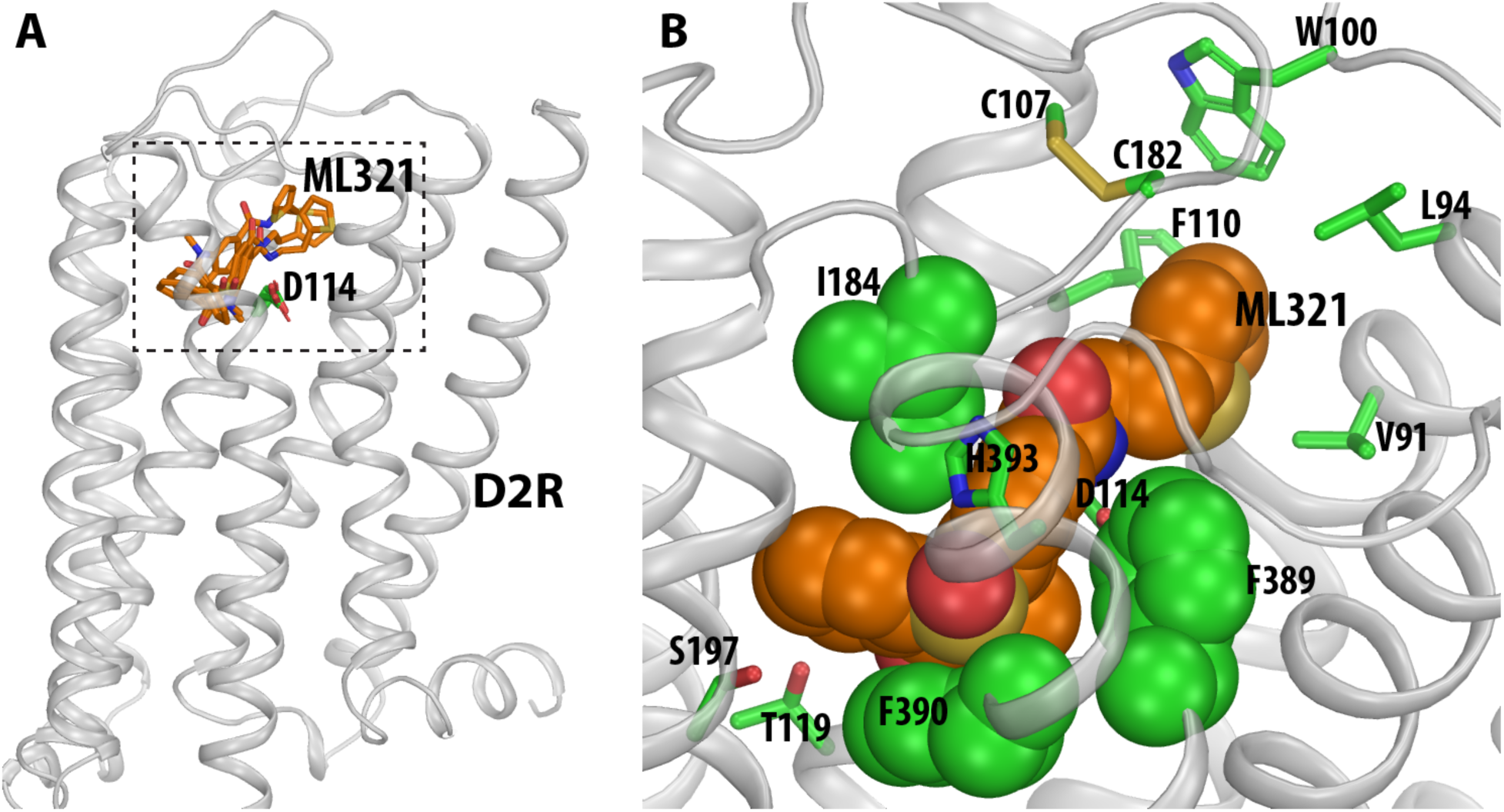
The computationally identified binding pose of ML321 at the D2R. (**A**) A side view of the D2R model in complex with several docked poses of ML321 (shown in orange sticks). The extracellular and intracellular sides of the receptor are on the top and bottom of the figure, respectively. The location of the ligand binding pocket is indicated by a dotted box. (**B**) A zoom-in view of the ligand binding pocket bound with the converged binding pose of ML321. The key contact residues are shown in green representations. Note the bulky hydrophobic or aromatic sidechains of Ile184^EL2.52^, Phe389^6.51^, and Phe390^6.52^ are tightly and complementarily packed with the dibenzothiazepine moiety of ML321, while the thiophene moiety protrudes into a sub-pocket formed by Val91^2.61^, Leu94^2.64^, Trp100^EL1.50^, Phe110^3.28^, and Cys182^EL2.50^.

Within the OBS, which is enclosed by transmembrane segment 3 (TM3), TM5, TM6, TM7, and extracellular loop 2 (EL2) ^30^, the concave side of the dibenzothiazepine moiety of ML321 clamps onto the hydrophobic sec-butyl sidechain of Ile184^EL2.52^ (superscripts denote Ballesteros-Weinstein numbering^29^, applied to extracellular loop regions as in Lane et al. ^36^), while the phenyl sidechains of Phe389^6.51^ and Phe390^6.52^ stack with the dibenzothiazepine moiety from its convex side (**Figure 5B)**. Such a complementary packing coordinates a hydrogen bond between the sidechain of His393^6.55^ and the sulfoxide oxygen of the dibenzothiazepine moiety. In addition, while ML321 does not possess a charged pyramidal nitrogen, its amide nitrogen can form a hydrogen bond with the sidechain carboxyl group of Asp114^3.32^. Parenthetically, this aspartate residue in TM3 is highly conserved within aminergic receptors and it typically forms a salt bridge with the protonated nitrogen found within most aminergic ligands^30^. Notably, in this pose, the dibenzothiazepine moiety does not form any direct polar interactions with the serine residues in TM5. Importantly, ML321 occupancy of the OBS as shown in **Figure 5B** is consistent with our observations (**Figure 3** and **Figure S2**) that ML321 exerts competitive antagonism at the D2R.

In the converged ML321 pose, the thiophene moiety protrudes into a SBP of the D2R formed by Val91^2.61^, Leu94^2.64^, Trp100^EL1.50^, Phe110^3.28^, and Cys182^EL2.50^, all of which interact with the thiophene moiety (**Figure 5B**). Notably, in our MD simulations, the thiophene ring pushes back and forth with the phenyl sidechain of Phe110^3.28^ (**Supplemental Video**). When this phenyl ring rotates outward towards the membrane, ML321 is in a more extended pose; when the phenyl ring faces inward, the thiophene ring is oriented away from the SBP resulting in a relatively folded configuration of ML321. This observation suggests that the bulky phenyl sidechain of Phe110^3.28^ may not favor the presence of the thiophene in this SBP.

To validate the computationally identified pose of ML321 in the D2R, we individually mutated the predicted binding site residues and assessed their effects on ML321 activity using radioligand binding competition and Go BRET functional assays as described in **Figure 1B** and **Figure S2B**, respectively. We initially performed saturation binding assays using [^3^H]-methylspiperone to assess the expression of the mutant receptors. These constructs expressed to a similar degree as the wild-type D2R (data not shown); however, Val91^2.61^A and Trp100^EL1.50^A were impaired in their binding to [^3^H]-methylspiperone thereby negating assessment of ML321 affinity using competition binding with the latter mutants. (A Val91^2.61^A mutant of the D2R had also previously been shown not to bind [^3^H]-methylspiperone^37^). Similarly, most of the mutant receptors exhibited dopamine-stimulated Go activation, except for Thr119^3.37^A and Ser197^5.46^, thus resulting in an inability to assess ML321 antagonism with those mutants. For the Go BRET assays, we stimulated the wild-type and mutant receptors with an EC_80_ concentration of dopamine, as determined for each mutant receptor, and assessed the ability of ML321 to inhibit the dopamine response. The EC_50_ and IC_50_ values for dopamine and ML321, respectively, were determined from their concentration-response curves and expressed as mutant/wild-type ratios as shown in **Table S3**. Similarly, [^3^H]-methylspiperone competition binding assays were performed using both ML321 and unlabeled spiperone (as a control) with the wild-type and the applicable mutant receptors. In each assay, we calculated the K_i_ values for the competing ligands and then expressed them as mutant/wild-type ratios to assess the impact of the mutations on the ligand binding affinities (**Table S4**).

In the OBS, the importance of the hydrogen bond between the nitrogen on the imidazole sidechain of His393^6.55^ and the sulfoxide of the dibenzothiazepine moiety is supported by the ∼10-fold reduction of ML321 binding affinity seen with H393^6.55^A (**Table S4**). Similarly, the reduced functional potencies of ML321 seen with Ile184^EL2.52^A and Phe390^6.52^A (∼2- and >40-fold, respectively) are consistent with their roles in shaping the OBS to accommodate the tricyclic dibenzothiazepine moiety, as well as forming favorable hydrophobic and aromatic interactions with it (**Table S3)**. Consistent with these results, the affinity of ML321 for binding to Phe390^6.52^A is reduced by 19-fold (**Table S4**). In our MD simulations, only Ser197^5.46^ among the three TM5 serines can potentially form an indirect polar interaction through a water molecule to the dibenzothiazepine moiety. However, its mutation, Ser197^5.46^A, did not have a significant effect on ML321 binding or function, suggesting that the polar decorations of the dibenzothiazepine moiety are unlikely to face TM5, as demonstrated by our binding pose of ML321 (**Figure 5B**).

In the SBP, Val91^2.61^A reduced the potency of ML321 by ∼5 fold for inhibiting dopamine-stimulated Go activation, while Trp100^EL1.50^A essentially eliminated ML321 activity, confirming the involvement of these residues in ML321 binding (**Table S3**). Similarly, Leu94^2.64^A reduced the functional potency of ML321 by ∼2-fold (**Table S3**) although this difference did not achieve statistical significance in the binding assay (**Table S4**). This might be due to Leu94^2.64^ being located on top of the SBP, where the hydrophobic mutation to Ala is more likely to be tolerated. Strikingly, Phe110^3.28^A improved both the binding affinity (∼20-fold) and the functional potency (∼5-fold) of ML321. This rarely observed gain-of-function mutation is consistent with the steric clash between the thiophene moiety of ML321 and the phenyl sidechain of Phe110^3.28^ in our binding pose (see above), whereby the removal of the bulky phenyl ring allows ML321 to fit more comfortably in an extended low-energy conformation. Overall, our pharmacological characterization of the binding site mutations strongly supports the computationally-identified ML321 binding pose at the D2R.

### ML321 Binding to the D2R is Sensitive to Na^+^

Previously, it was determined that butyrophenone antagonists, such as spiperone and risperidone, protrude deep into the OBS of the D2R and occupy a sub-pocket defined by Ile^3.40 36^, which is located near the receptor’s conserved Na^+^ binding site at Asp^2.50 38–40^. Lane and colleagues^36^ have proposed that interactions with this Ile^3.40^ sub-pocket render butyrophenone binding to the D2R insensitive to Na^+^. In contrast, Na^+^ is known to enhance the binding of substituted benzamide antagonists, such as sulpiride and eticlopride, that lack interactions with the Ile^3.40^ sub-pocket of the D2R^36, 41^. As our modeling data suggest that ML321 does not occupy the Ile^3.40^ sub-pocket, we were interested in determining whether the binding of ML321 to the D2R would be modulated by Na^+^. **Figure 6** illustrates radioligand binding competition experiments conducted using Tris buffer in the absence or presence of 140 mM NaCl. **Figure 6A** clearly shows that inclusion of NaCl in the buffer enhances the affinity of ML321 for the D2R (K_i_ = 72 nM with Na^+^ *versus* >10 μM without Na^+^). We also examined the Na^+^ sensitivity of ML321 binding to the D3R despite its lower affinity for this receptor subtype. **Figure 6B** shows that ML321’s affinity for the D3R may be enhanced by Na^+^; however, the low affinity of ML321 for the D3R made it difficult to quantify this effect. Overall, these results agree with and support our modeling data showing that ML321 does not occupy the Ile^3.40^ sub-pocket of the D2R and they further demonstrate that ML321 binding to the D2R is enhanced by Na^+^.

**Figure 6.**
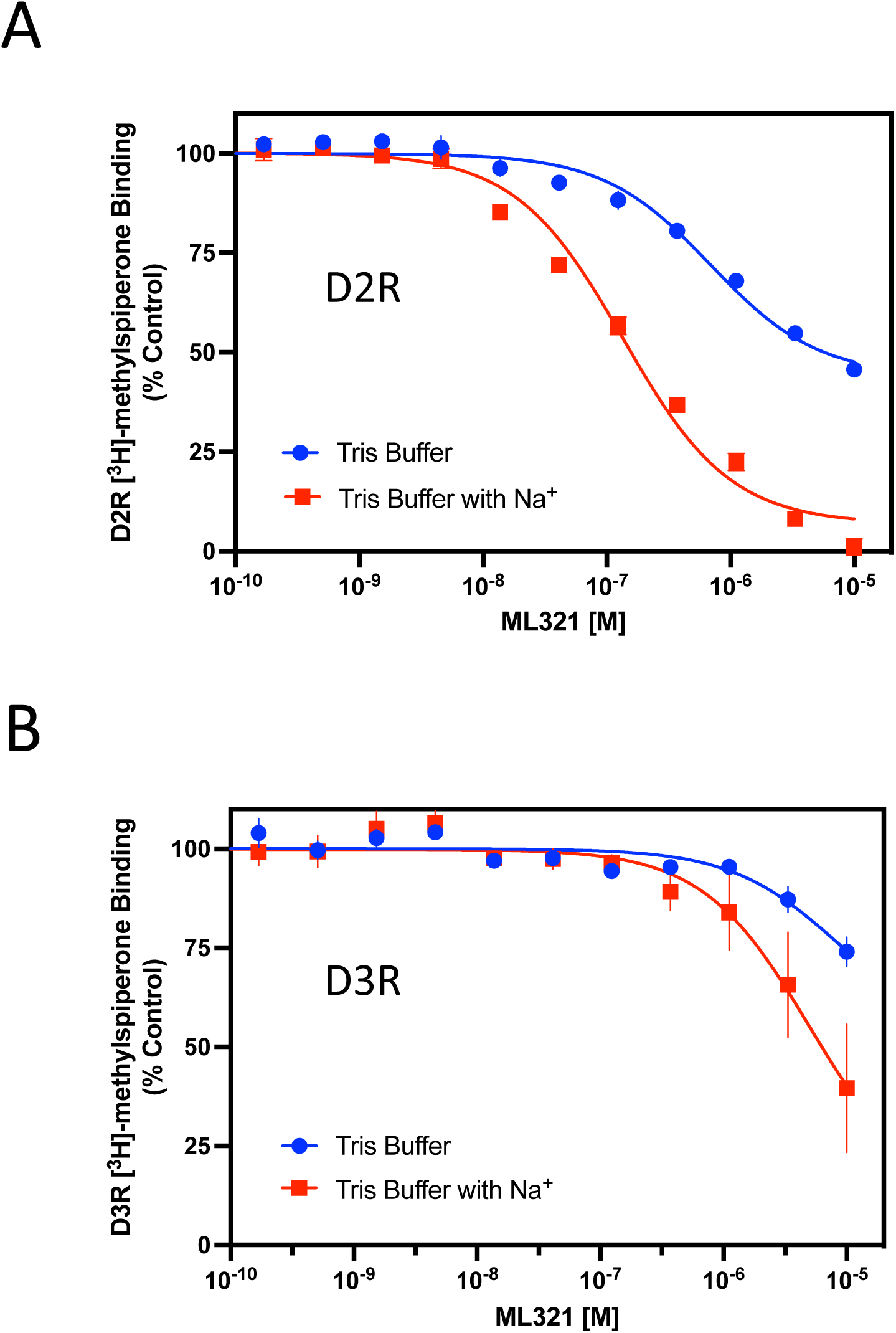
The binding of ML321 to the D2R is regulated by Na^+^. D2R and D3R radioligand binding assays were performed as described in the Methods using Tris buffer in the absence or presence of 140 mM NaCl. The data are expressed as percentage of the control specific binding and represent mean ± SEM values from three independent experiments each performed in triplicate. Mean ML321 Ki values [95% C.I.] for each receptor were calculated from the IC_50_ values using the Cheng-Prusoff equation^31^. (**A**) D2R competition binding curves ± NaCl. In the presence of Na^+^, the mean ML321 Ki = 72.5 nM [56.3-93.3] whereas in the absence of Na^+^ the Ki >10 μM. (**B**) D3R competition binding curves ± NaCl. In the absence or presence of Na^+^, the ML321 Ki values are >10 μM.

### PET Imaging Demonstrates that ML321 Penetrates the CNS and Occupies the D2R in a Dose-Dependent Manner

We previously demonstrated that ML321 is brain penetrant in rodents (brain:plasma partition coefficient = 0.2) ^28^; however, we also wished to establish that it could effectively engage the D2R within the CNS. To demonstrate this point, we conducted a micro-PET imaging study in rhesus monkeys using the PET tracer [^11^C]SV-III-130 that has previously been employed to image the D2R in the CNS^42, 43^. [^11^C]SV-III-130 exhibits a K_i_ of 0.22 nM for the D2R ^44^ (SV-III-130 = compound **6**). **Figure 7** shows an experiment in which [^11^C]SV-III-130 is administered (i.v.) to rhesus monkeys and the subsequent uptake of the tracer into the caudate, putamen, and cerebellar regions of the brain is continuously monitored up to 100 min. As shown, there is rapid uptake of [^11^C]SV-III-130 into all three brain regions, although the level of [^11^C]SV-III-130 rapidly declines in the cerebellum, a brain region that is nearly devoid of D2R expression. In contrast, the uptake of [^11^C]SV-III-130 in the caudate and putamen, brain regions of very high D2R expression, is maintained throughout the experiment (**Figure 7**, see baseline imaging). However, if either 1 mg/kg or 5 mg/kg of ML321 is administered 20 min after [^11^C]SV-III-130 infusion, there is a significant decline of [^11^C]SV-III-130 uptake in both the caudate and putamen due to its displacement by ML321 at the D2R. Moreover, the effects of ML321 on [^11^C]SV-III-130 levels occur in a dose-dependent fashion (**Figure 7**). Notably, ML321 had no effect on the rapid decline of [^11^C]SV-III-130 in the cerebellum. Similar results were previously observed with the D2-like receptor antagonist eticlopride with respect to displacing [^11^C]SV-III-130 uptake in the brain^42^. Taken together, these results confirm that ML321 is brain-penetrant and they further show that it can occupy the D2R in relevant brain regions in a dose-dependent manner.

**Figure 7.**
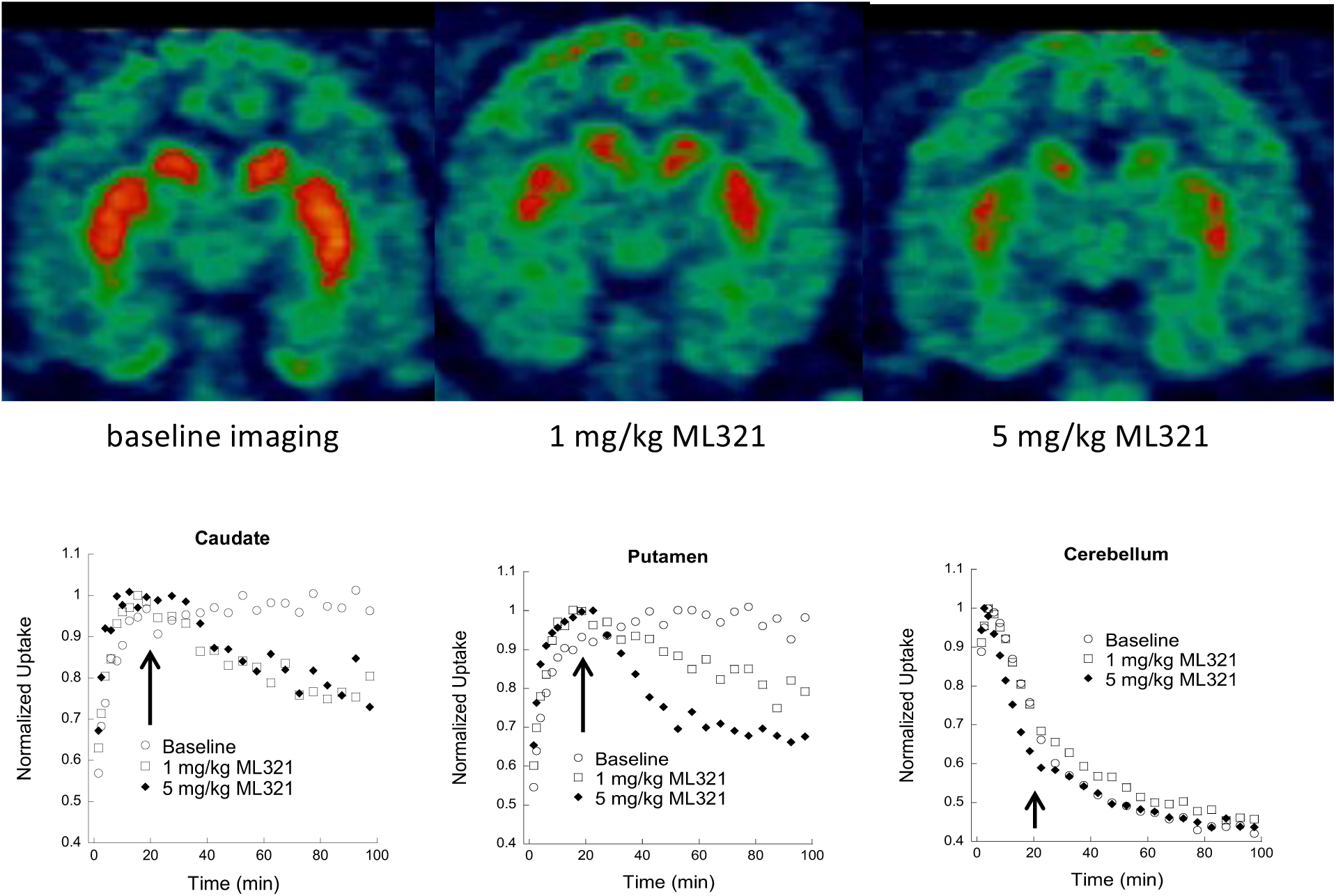
Displacement of the micro-PET tracer [^11^C]SV-III-130 by ML321 in brains of non-human primates. MicroPET imaging studies were conducted as described in the Methods. Uptake of the radioactive micro-PET tracer [^11^C]SV-III-130 was monitored continuously for 100 min after injection (i.v.). For the drug treatments, either 1 or 5 mg/kg of ML321 was administered (i.v.) 20 min (arrows) after tracer infusion. **Top:** coronal brain sections are shown illustrating the uptake of [^11^C]SV-III-130 into the caudate and putamen (orange areas) under baseline conditions (*left*) or after the administration of 1 mg/kg (*middle*) or 5 mg/kg (*right*) of ML321 (t = 100 min). **Bottom:** tissue time-activity curves illustrating the uptake of [^11^C]SV-III-130 into the caudate (*left*), putamen (*middle*), or cerebellum (*right*). The data are normalized to the maximum uptake seen in each brain region. After 20 min (arrows), the animals were treated with either vehicle or the indicated doses of ML321. The representative results from single experiments are shown, which were performed three times with similar results.

### ML321 Selectively Antagonizes the D2R vs. the D3R Using *In Vivo* Paradigms

Previously we showed that following administration (i.p.) to mice, ML321 achieved maximum plasma and brain concentrations within 15 min (brain Cmax/plasma Cmax = 0.2) and exhibited t_/1/2_ values of 1.67 and 1.32 hr in plasma and brain, respectively^28^. We deemed these results sufficient to conduct behavioral experiments. Initially, we sought to determine if we could recapitulate the D2R > D3R selectivity of ML321 that we observed *in vitro* using *in vivo* behavioral models. To accomplish this, we used agonist-stimulated hypothermia and yawning responses in rats, which are mediated by the D2R and D3R, respectively^45^. **Figure 8A** shows the changes in body temperature from treating rats with 1.0 mg/kg of the D2R-preferring agonist sumamirole, administered subcutaneously. As observed previously^45^, treatment with sumanirole alone promoted a decrease (>1°C) in body temperature. Notably, the hypothermia produced by sumanirole was significantly antagonized by ML321 using doses of 3.2 or 10 mg/kg (*p*-values <0.001). In contrast, treatment with 10 mg/kg of ML321 alone appeared to have no effect on body temperature.

**Figure 8.**
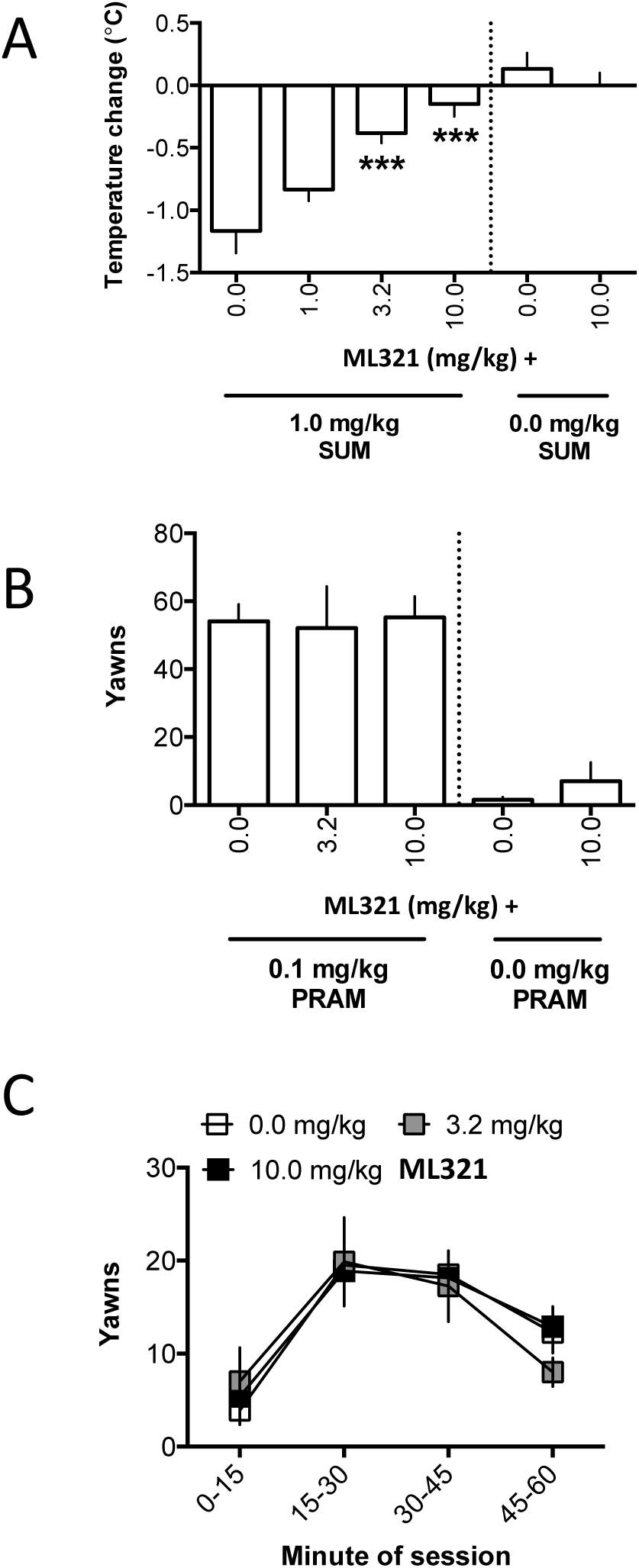
Effects of ML321 on D2R-mediated hypothermia or D3R-mediated yawning in rats. Measurements of hypothermia and yawning were performed as described in the Methods. All data are presented as the mean ± SEM. **(A)** Measurement of hypothermia: change in rats’ body temperature following injection (s.c) with vehicle (0.0 mg/kg) or 1.0, 3.2, or 10.0 mg/kg ML321 and 1.0 mg/kg sumanirole (SUM). Experimental groups contained 6 animals each except for the 10.0 mg/kg ML321 + vehicle group, which contained 3 animals. For the ML321 + sumanirole groups, following a significant effect of ML321 dose in the one-way ANOVA [*F*(3,20) = 14.69, *p* < 0.001], pairwise comparisons to vehicle were made *post hoc* using Dunnett’s tests (two-tailed): \*\*\**p* < 0.001, significantly different from 0.0 mg/kg ML321 (vehicle) + 1.0 mg/kg sumanirole. For the 10 mg/kg ML321 + vehicle group, the data are presented for reference, but were not statistically analyzed given the small number of animals tested. **(B)** Measurement of yawning: number of yawns made in 60 min following injection (s.c.) with vehicle (0.0 mg/kg) or 3.2 or 10.0 mg/kg ML321 and 0.1 mg/kg pramipexole (PRAM). Experimental groups contained 8 animals except for the 10.0 mg/kg ML321 + vehicle group, which contained 3 animals. For the ML321 + pramipexole groups, the one-way ANOVA for dose was not significant. [*F*(2,21) = 0.03, *p* = 0.96]. For the 10 mg/kg ML321 + vehicle group, the data are presented for reference, but were not statistically analyzed given the small number of animals tested. **(C)** To obtain a time-course of pramipexole-induced yawning, the 60 min observation session in **B** was divided into 4 blocks of 15 min each, and the data were reanalyzed as the mean yawns in each time-block. A two-way ANOVA detected a significant effect of block [F(3,84)=16.54, *p* <0.001]; however, the mean yawns/block were not affected by ML321 dose (non-significant main effect and interaction).

**Figure 8B** shows the total yawns made by rats during a 60 min observation session following pretreatment with various doses of ML321 and treatment with 0.1 mg/kg of the D3R-preferring agonist pramipexole, administered subcutaneously. Notably, yawning following treatment with 0.1 mg/kg pramipexole was not significantly affected by any ML321 pretreatment dose. In addition, treating rats with 10 mg/kg ML321 alone did not appear to promote yawning. To determine that ML321 did not exhibit a transient effect (i.e., a slow onset or rapid offset of action) that was obscured when the total number of yawns was analyzed, the time-course of yawning during the 60-min session was analyzed by binning the data into 15 min blocks (**Figure 8C**) across the 60-min session. Yawning significantly changed over the course of the session (*p*<0.001); however, neither the main effect of ML321 dose nor the block-by-dose interaction was significant. Taken together, these data show that a dose of ML321 can be chosen to selectively attenuate a D2R-mediated response (hypothermia) while exerting no effect on a D3R-mediated response (yawning). Thus, ML321 can be used to selectively antagonize the D2R *in vivo*.

### ML321 is Active in Animal models that Predict Antipsychotic Activity, yet it does not Promote Catalepsy

We were interested in determining if ML321 is effective in animal models that are predictive of antipsychotic activity in humans. Amphetamine (AMPH)- and phencyclidine (PCP)-induced hyperlocomotion are two pharmacological models that are commonly used^46^. The AMPH-induced response is dependent on striatal dopamine release, whereas the behavioral effects of PCP are thought to be mediated by cortical disinhibition and activation of the corticostriatal pathway^46^. As antipsychotics are D2R antagonists or low-efficacy partial agonists, they have been shown to attenuate AMPH- and PCP-induced hyper-locomotor responses. In the experiment in **Figure 9**, mice were first injected (i.p.) with vehicle or various doses of ML321 and placed in an open field for 30 min while monitoring locomotion. After 30 min, the mice were removed and administered (i.p.) either vehicle (**Figure 9A**), 3 mg/kg AMPH (**Figure 9 B, D**), or 6 mg/kg PCP (**Figure 9 C, E**) and returned to the open field for 90 min. No significant baseline group differences were observed and, as expected, habituation of locomotor activity across baseline was observed (**Figure 9A-C**). While the kinetics of the AMPH- and PCP-induced responses were robust, the ML321 dose-dependent reductions in activity are clearly evident when the data are presented as cumulative locomotion. Notably, when tested alone at the highest dose (5 mg/kg ML321-vehicle), no significant effects on locomotor activity were identified as compared to the vehicle-vehicle control (**Figure 9D-E)**. Thus, at this dose ML321 alone does not have locomotor stimulating or inhibiting effects in mice. However, ML321 depressed the vehicle-AMPH induced hyperlocomotion in a dose-dependent manner with 1 and 5 mg/kg ML321-AMPH (*p*-values ≤ 0.002), and a similar relationship in suppression was observed by reducing the vehicle-PCP stimulated activity with 0.25, 0.5, 1, and 5 mg/kg ML321-PCP (*p*-values ≤ 0.025). Importantly, the highest dose, 5 mg/kg ML321-AMPH or ML321-PCP, completely blocked both the AMPH- and PCP-induced responses such that these values were not significantly different from either the vehicle-vehicle or 5 mg/kg ML321-vehicle controls. Notably, the ML321 dose-response relationships observed in this study approximate those exhibited in the experiments shown in **Figures 7 and 8**. Further, they predict that ML321 would exhibit antipsychotic efficacy in schizophrenic patients.

**Figure 9.**
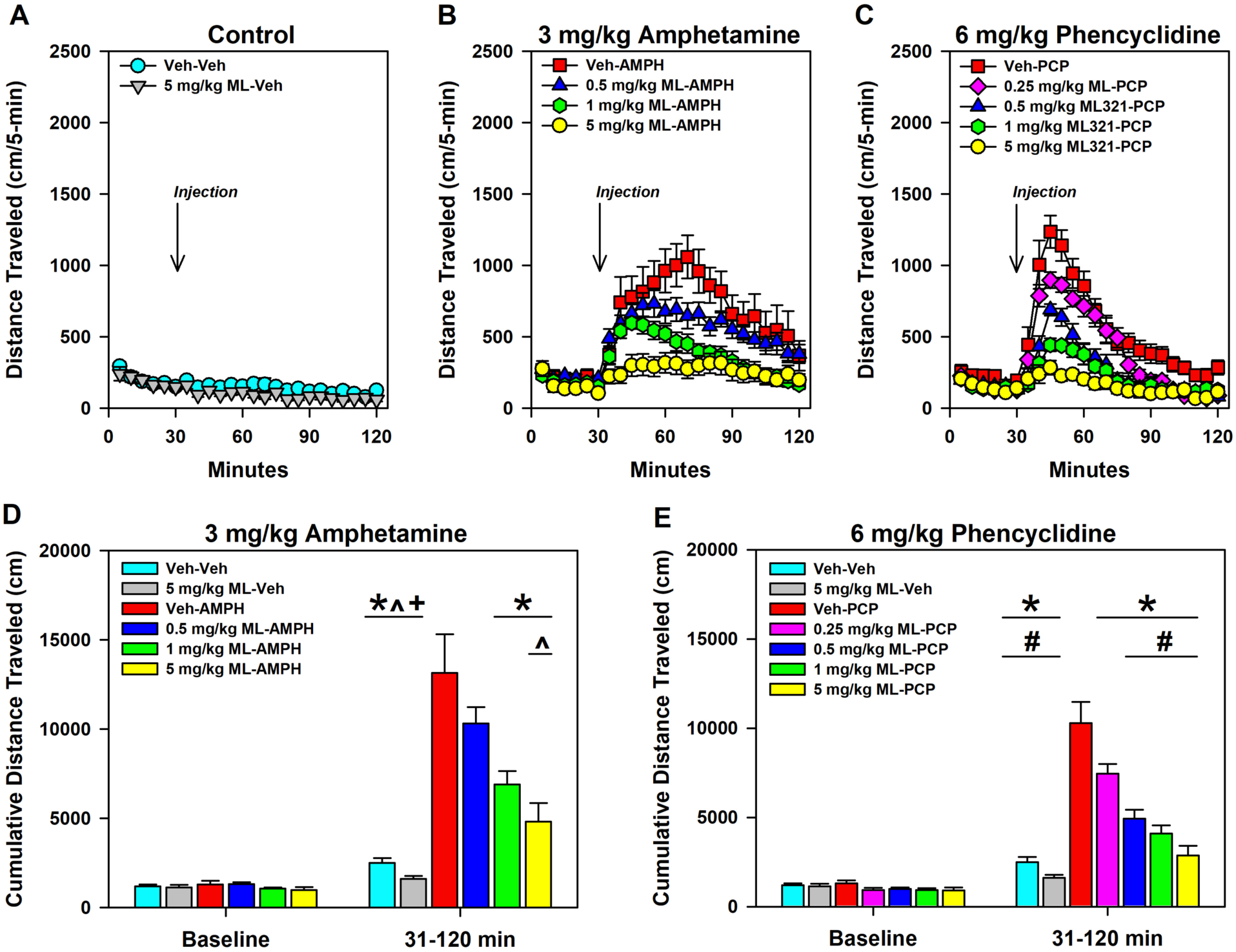
ML321 displays potent antipsychotic-like activity in a hyperlocomotion study. C57BL/6J mice were injected (i.p) with either the vehicle (Veh) or the indicated doses of ML321 (ML) and placed in an open field with assessment of locomotor activity as described in Methods. After 30 min, the mice were removed and injected (i.p.) with either Veh, 3 mg/kg of amphetamine (AMPH), or 6 mg/kg phencyclidine (PCP) and immediately returned to the open field for a further 90 min. Locomotor activities (distance traveled) are shown as 5-min binned intervals **(A-C)** or as cumulative locomotion **(D-E)**. For the AMPH experiment **(B, D)**, a RMANOVA for baseline (0-30 min) identified a significant effect of time [F(5,285)=25.168, *p*<0.001], while an analysis of the post-injection interval (31-120 min) revealed the time [F(17,969)=20.532, *p*<0.001], treatment [F(5,57)=18.231, *p*<0.001], and time by treatment interaction [F(85,969)=3.860, *p*<0.001] were significant. For the PCP study **(C, E)**, a RMANOVA for baseline (0-30 min) observed a significant effect of time [F(5,335)=28.707, *p*<0.001], whereas an analysis of the stimulated interval (31-120 min) reported the time [F(17,1139)=102.322, *p*<0.001], treatment [F(6,67)=29.915, *p*<0.001], and the time by treatment interaction [F(102,1139)=12.401, *p*<0.001] to be significant. **(D-E)** A one-way ANOVA found baseline (0-30 min) locomotor activities were not significantly different between the Veh-Veh and 5 mg/kg ML-Veh groups. However, following administration of the psychostimulants (31-120 min), one-way ANOVAs demonstrated significant stimulated effects for the AMPH [F(5,62)=18.231, *p*<0.001] and PCP [F(6,73)=26.915, *p*<0.001] experiments. All data are presented as means ± SEMs. N=11 for the 0.25 ML-PCP group, N=13 for the Veh-Veh control, and N=10 for all other groups. \**p*<0.05, Veh-AMPH or Veh-PCP *vs.* the other indicated groups; ^#^*p*<0.05, 0.25 mg/kg ML-PCP *vs.* the Veh-Veh, 5 mg/kg ML-Veh, and the 0.5-5 mg/kg ML-PCP groups; ^^^*p*<0.05, 0.5 mg/kg ML-AMPH *vs.* the 5 mg/kg ML-AMPH group; ^+^*p*<0.05, 1 mg/kg ML-AMPH *vs.* the Veh-Veh and 5 mg/kg ML-Veh groups.

In a second study, we examined pre-pulse inhibition (PPI) of acoustic startle in mice following AMPH or PCP administration. PPI of the acoustic startle reflex refers to the ability of a weak acoustic stimulus preceding a stronger startling stimulus to inhibit the response to the second stimulus. PPI is considered a form of sensorimotor gating since it refers to the ability of a sensory event to suppress a motor response. PPI deficits have been documented in schizophrenic patients ^47^ and they can be induced in animals with psychotomimetics such as AMPH or PCP, and these deficits can be reversed by antipsychotic drugs^48^. In the study shown in **Figure 10**, mice were injected (i.p.) with either vehicle or various doses of ML321. Ten minutes later, they were given the vehicle, 3 mg/kg AMPH, or 6 mg/kg PCP, and were acclimated to a 65 dB white-noise background for 10 min in the PPI apparatus. After 5 min, mice were presented with combinations of startle (120 dB), pre-pulse-pulse (4, 8, and 12 dB over the 65 dB background followed by 120 dB), and null trials over 25 min. For the AMPH study, null activity was increased in the vehicle-AMPH mice relative to the vehicle-vehicle, 5 mg/kg ML321-vehicle, 0.25 mg/kg ML321-AMPH, and the 5 mg/kg ML321-AMPH groups (*p*-values ≤ 0.035) **(Figure S4A)**. Activity was also enhanced in the 0.5 and 1 mg/kg ML321-AMPH animals compared to the vehicle-vehicle control and the 5 mg/kg ML321-AMPH group. When startle activity was examined, responses were higher in the vehicle-vehicle control than the 5 mg/kg ML321-vehicle and the 0.25, 0.5, and 1 mg/kg ML321-AMPH groups (*p-*values ≤ 0.015) **(Figure S4B)**. In the PCP experiment, all doses of ML321 with PCP increased null activity compared to the vehicle-vehicle and 5 mg/kg ML321-vehicle controls (*p*-values ≤ 0.016) **(Figure S4C)**, whereas startle reactivity was potentiated in the vehicle-vehicle groups relative to the 0.25 mg/kg ML321-PCP mice (*p* = 0.045) **(Figure S4D)**. When PPI was analyzed with 5 mg/kg ML321-vehicle, no significant effects were observed compared to the vehicle-vehicle in either the AMPH or PCP investigations **(Figure 10)**. Hence, ML321 alone does not disrupt PPI. By contrast, both AMPH and PCP significantly disrupted PPI compared to this control (*p*-values<0.001), whereas ML321 dose-dependently reversed the AMPH- and PCP-induced PPI suppression with the 5 mg/kg ML321-AMPH or ML321-PCP treatments normalizing PPI to baseline levels (**Figure 10**). Taken together, these results further predict that ML321 has a high potential for antipsychotic activity in schizophrenic patients.

**Figure 10.**
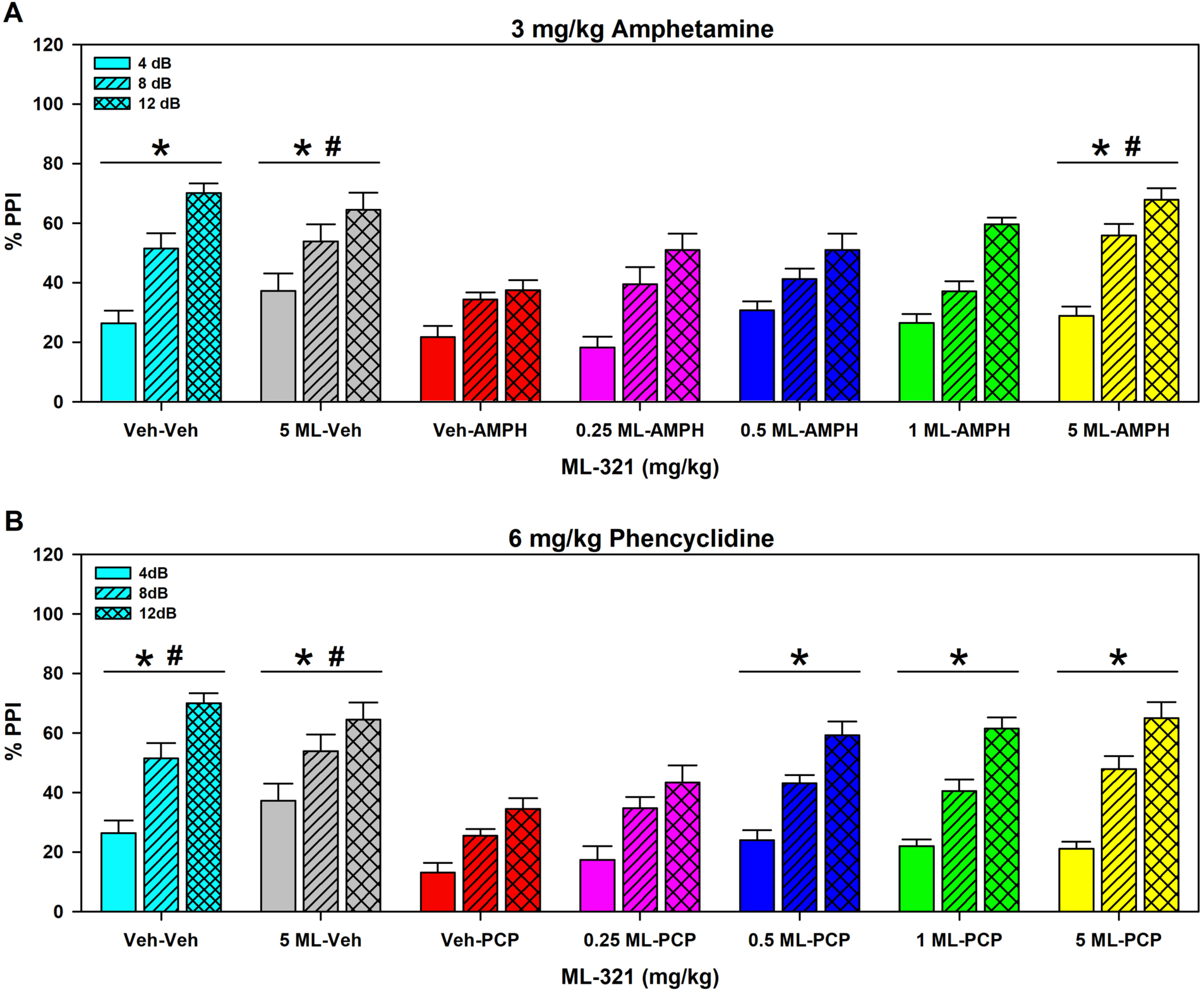
ML321 reverses psychostimulant-induced impairments in pre-pulse inhibition of the acoustic startle. C57BL/6J mice were administered (i.p.) vehicle (Veh) or 0.25, 0.5, 1.0, or 5 mg/kg doses of ML321 (ML) in their home-cages. Ten minutes later, animals were given the Veh, 3 mg/kg amphetamine (AMPH) **(A)**, or 6 mg/kg phencyclidine (PCP) **(B)** and acclimated to a 65 dB white-noise background for 10 min in the PPI apparatus. After 5 min, mice were presented with combinations of startle (120 dB), pre-pulse-pulse (4, 8, and 12 dB over the 65 dB background followed by 120 dB), and null trials over 25 min (see Methods). Activity was recorded during all trials. The data are presented as % PPI = [1 – (pre-pulse trials/startle-only trials)] x 100. A RMANOVA for the AMPH study noted significant effects of PPI [F(2,122)=147.032, *p*<0.001], treatment [F(6,61)=6.120, *p*<0.001], and the PPI by treatment interaction [F(12,122)=2.366, *p*=0.009]. A RMANOVA for the PCP experiment determined that the effects of PPI [F(2,122)=176.947, *p*<0.001], treatment [F(6,61)=8.393, *p*<0.001], and the PPI by treatment interaction [F(12,122)=1.883, *p*=0.043] were significant. The data are presented as means ± SEMs. N=9 mice for the vehicle-vehicle, vehicle-AMPH, and vehicle-PCP groups; N=10 mice for all other groups. \**p*<0.05, for the AMPH study: Veh-AMPH *vs.* the Veh-Veh, 5 mg/kg ML-Veh, and the 5 mg/kg ML-AMPH groups; or \**p*<0.05, for the PCP experiment: Veh-PCP *vs.* the Veh-Veh, 5 mg/kg ML-Veh, and the 0.5 to 5 mg/kg ML-PCP groups; ^#^*p*<0.05, for the AMPH investigation: 0.25 mg/kg ML-AMPH *vs.* the 5 mg/kg ML-Veh and the 5 mg/kg 5 ML-AMPH groups; or for the PCP study: ^#^*p*<0.05, 0.25 mg/kg ML-PCP *vs.* the Veh-Veh and the 5 mg/kg ML-Veh groups.

Most antipsychotics are associated with parkinsonian-like extrapyramidal symptoms (EPS) that involve bradykinesia and/or tremors, which are frequently, but not always, observed with high and/or prolonged dosing, as it represents an on-target side effect. While the mechanism(s) underlying EPS is uncertain (see below), this can be modeled in rodents through examining drug-produced catalepsy, which is characterized as postural rigidity. We initially examined ML321’s liability to produce catalepsy in mice compared to the typical antipsychotic haloperidol (HAL) using the horizontal rod test (**Figure 11**). Mice were injected (i.p.) with vehicle or various doses of HAL or ML321 and were evaluated for catalepsy 60 min later. Compared with baseline (no drug), HAL produced significant catalepsy at 1 and 10 mg/kg (*p*-values<0.001). Additionally, the cataleptic responses to 1 and 10 mg/kg HAL were significantly different from each other (*p*<0.001) and both were higher than those for the lower doses of HAL (*p*-values<0.001). By striking contrast, ML321 produced very little catalepsy even when evaluated using a high 10 mg/kg dose, which is at or above a maximal dose for producing ML321-mediated effects in the other behavioral assays (**Figures 8-10**). To confirm our finding that ML321 is relatively ineffective in producing catalepsy, we also used the inclined screen test for catalepsy (**Figure S5**). In this experiment, the mice were injected (i.p.) with vehicle or 10 mg/kg ML321 or 10 mg/kg HAL and tested for catalepsy after 60 min. As was observed with the horizontal bar test, HAL induced significant catalepsy according to both test indices (*p*-values<0.001) whereas ML321 was ineffective. Taken together, these results suggest that ML321 may exhibit very little EPS in patients, thus behaving as an “atypical” antipsychotic.

**Figure 11.**
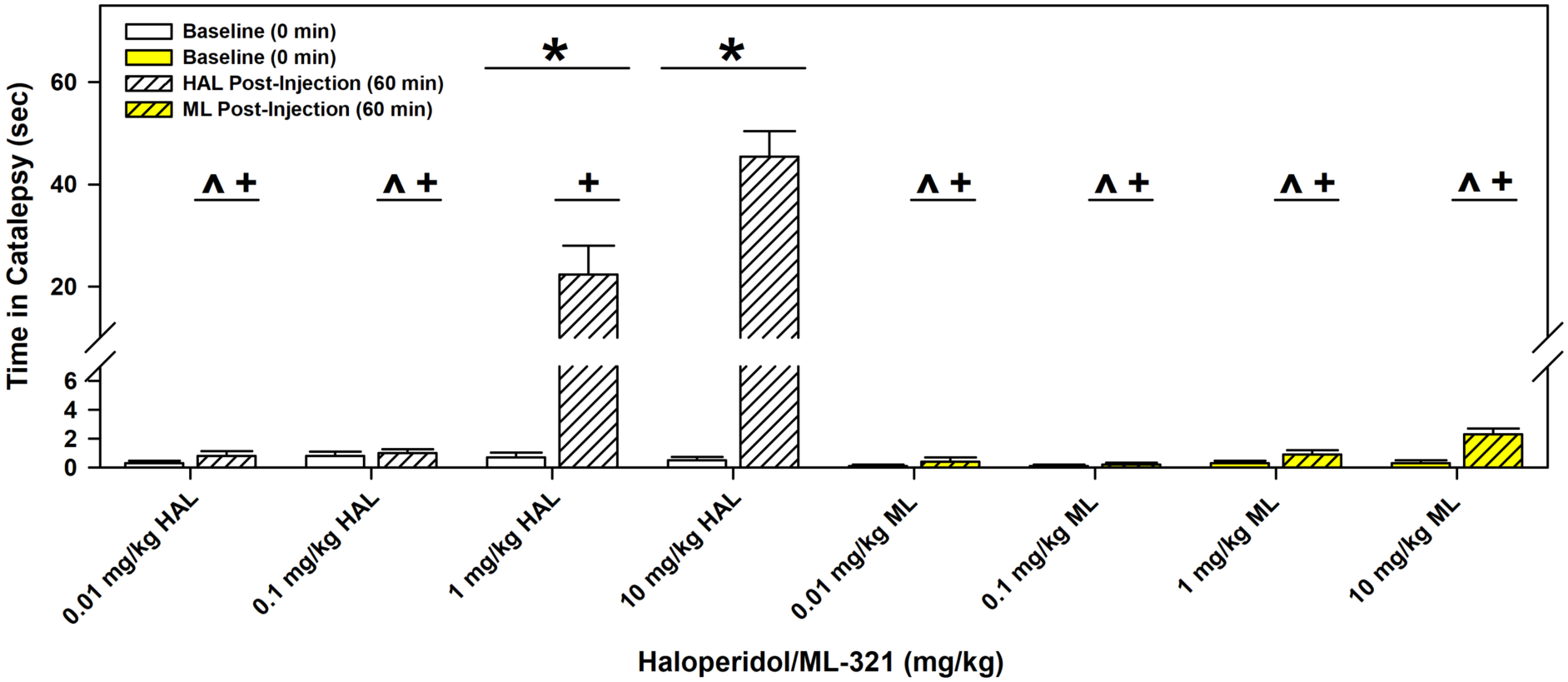
ML321 has low cataleptic potential in C57BL/6J mice compared to haloperidol in the horizontal bar test. Baseline responses were recorded and then separate groups of C57BL/6J mice were injected with either vehicle (Veh) or the indicated doses of haloperidol (HAL) or ML321 (ML). Mice were tested 60 min later for catalepsy. The latency for a mouse to remove its paws from the bar was recorded as an index of catalepsy (60 second maximum time). A RMANOVA observed the main effects of time [F(1,72)=85.874, *p*<0.001] and treatment [F(7,72) =35.542, *p*<0.001], as well as the time by treatment interaction [F(7,72)=37.267, *p*<0.001] to be significant. The data are presented as means ± SEMs. N=10 mice/group. \**p*<0.05, baseline *vs.* 60 min; ^^^*p*<0.05, 1 mg/kg HAL *vs.* 0.01 and 0.1 mg/kg HAL, and all doses of ML; ^+^*p*<0.05, 10 mg/kg HAL *vs.* 0.01 and 0.1 mg/kg HAL, and all doses of ML.

### Kinetic Studies Reveal that ML321 Exhibits Slow Association and Fast Dissociation Rates of Binding to the D2R

Drug-induced catalepsy in rodents is highly correlated with EPS in humans, and “typical” antipsychotics, like HAL promote both. As noted above, “atypical” antipsychotics promote less EPS and, similarly, they produce less catalepsy in rodents^49^. Multiple theories have been suggested for the decreased EPS observed with atypical drugs with a prominent one being that EPS is correlated with the kinetics of drug binding to the D2R. Approximately 20 years ago, Kapur and Seeman (2001) ^19^ initially posited that less EPS is associated with faster dissociation of an antipsychotic from the D2R (the “fast-off” hypothesis). More recently, Sykes et al. (2017) ^20^ have shown there is a high correlation of EPS with the association rate of antipsychotic drug binding such that a slower association rate, mainly reflected in “re-binding” to the receptor after dissociation, is correlated with less EPS (the “slow-on” hypothesis). In both hypotheses, the kinetics of drug binding reflect antipsychotic residence times at the receptor binding site such that shorter residence times permit some degree of dopamine occupancy of the D2R, and its associated signaling, leading to less EPS. Thus, antipsychotics with both fast-off and slow-on receptor binding kinetics would seemingly exhibit the least EPS. As such, we sought to determine the association and dissociation kinetics of ML321 to the D2R using the methods of Sykes et al. (2017) ^20^. Notably, we also used clozapine as a reference agent, as it is an atypical antipsychotic that produces little catalepsy in rodents or EPS in humans^10^ and it exhibits slow association and fast dissociation kinetics at the D2R ^20^. The atypicality of clozapine was once hypothesized to involve 5-HT_2A_ receptor antagonism^10, 18^; however, ML321 exhibits little affinity for the 5-HT_2A_ receptor (see above).

**Table 1** shows the results from a time-resolved fluorescence resonance energy transfer (TR-FRET) assay to measure association (*k*_on_) and dissociation (*k*_off_) rates as well as dissociation t_½_ values for clozapine and ML321 binding to the D2R. Interestingly, the association and dissociation kinetics for clozapine and ML321 are very similar to each other and the K_d_ value for ML321 derived from the kinetic data is essentially identical to the experimentally determined K_i_ value from competition binding experiments (see **Figure 1**). Similarly, the kinetic data for clozapine highly approximates those obtained by Sykes et al. (2017) ^20^, and the kinetically-derived K_d_ for clozapine agrees well with its published value^10, 50^. These results suggest that ML321 exhibits slow association and fast dissociation kinetics at the D2R relative to most antipsychotic drugs in current use. Further, they may explain why ML321 exhibits such a low propensity to produce catalepsy in the rodent models that we have employed.

**Table 1.** Association rates, dissociation rates, and dissociation t½ values of clozapine and ML321 binding to the D2R^*a*^.

| | $k_{\text{off}}$ (min<sup>-1</sup>) | $k_{\text{on}}$ (M<sup>-1</sup> min<sup>-1</sup>) | $t_{1/2}$ (min) | $K_d$ (nM) |
| --- | --- | --- | --- | --- |
| Clozapine | 1.6 ± 0.23 | 4.6 ± 0.7 X 10<sup>7</sup> | 0.47 ± 0.06 | 34.8 ± 0.06 |
| ML321 | 1.3 ± 0.22 | 2.3 ± 0.4 x 10<sup>7</sup> | 0.60 ± 0.07 | 56.5 ± 0.23 |

<sup>a</sup>Time-resolved fluorescence resonance energy transfer assays were used to measure association ( $k_{\text{on}}$ ) and dissociation ( $k_{\text{off}}$ ) rates and dissociation $t_{1/2}$ values as described in the Methods. Data are the means ± SEM of five (clozapine) or six (ML321) independent experiments performed in triplicate.

### Preliminary Toxicology/Safety Assessment

We conducted several studies (see Methods) to assess the potential toxicity of the ML321 scaffold. To evaluate the ability of ML321 to inhibit the hERG K^+^ channel, which can result in cardiotoxicity, we used HEK293 cells expressing the hERG channel and conducted whole cell voltage clamping experiments. ML321 was found to be ineffective in inhibiting the hERG channel with an estimated IC_50_ >25 μM (the highest concentration tested). We also performed a cytotoxicity screening panel that involved incubating HepG2 cells with various concentrations of ML321 for 72 hr and then measuring several cell-health parameters including cell count, nuclear size, DNA structure, cell membrane permeability, mitochondrial mass, mitochondrial membrane potential, and cytochrome c release. The AC_50_ values for ML321 having any effects on these parameters ranged from between 50 μM to >100 μM (the highest concentration tested). We also conducted Ames mutagenicity screening and found that ML321 was negative in this assay when tested up to 300 μM. Thus, while more extensive *in vivo* toxicity experiments involving the ML321 scaffold are required, these preliminary toxicity studies do not raise any immediate concerns.

## CONCLUSIONS

ML321 is a novel D2R antagonist with unprecedented selectivity *versus* other GPCRs. This attribute will allow it to serve as an excellent *in vivo* chemical probe or potentially as a therapeutic with drastically reduced off-target side effects. ML321 also exhibits unique receptor binding kinetics falling into the slow-on and fast-off category of ligands similar to that of the atypical antipsychotic clozapine. This observation and the lack of catalepsy observed in rodents upon ML321 administration suggest that this drug lead may function as an atypical antipsychotic with reduced on-target EPS. To-date, no other D2R antagonist has been shown to exhibit both of these pharmacological characteristics – exceptional receptor selectivity and slow-on/fast-off kinetics. Notably, however, ML321 itself may be metabolized too rapidly to serve directly as a therapeutic and will need to be further optimized into a derivative that exhibits a longer *in vivo* half-life. We have identified the metabolites of ML321 (unpublished observations) and are currently in the process of chemically optimizing this scaffold to create an advanced drug lead with improved pharmacokinetics for IND-enabling studies.

As noted above, D2R antagonism is effective for treating the positive symptoms of schizophrenia, but less efficacious for treating the negative or cognitive symptoms of this illness. While other drug targets are being investigated, particularly for the treatment of negative and cognitive symptoms^12–15^, it may prove difficult to develop a single drug that has the appropriate affinity and activity/efficacy for all, including yet unknown, targets needed to treat the entirety of schizophrenic symptomatology. Thus, similar to many other diseases and disorders, a combinatorial approach (administering two or more drugs, perhaps formulated together) may become the standard of care for treating schizophrenia (e.g., see^17^), and it is likely that one of these drugs will be a D2R antagonist (or low efficacy partial agonist).

In summary, we believe that the ML321 scaffold will be differentiated from other D2R antagonists used therapeutically by two unique characteristics: 1) an exceptional target selectivity such that off-target side effects are predicted to be very low or non-existent; and 2) unique D2R binding kinetics such that on-target side effects are predicted to be greatly reduced or absent. Such a drug will represent a significant advance in the treatment of schizophrenia and related disorders.

## MATERIALS AND METHODS

### Chemicals and Reagents

Coelenterazine-H and Coelenterazine 400a were purchased from Nanolight^TM^ Technology. Mutant receptor constructs were prepared by Bioinnovatise (Rockville, MD) in pcDNA3.1 vectors and inserts were verified by sequencing. All tissue culture media and supplies were obtained from ThermoFisher Scientific (Carlsbad, CA). All other compounds and chemicals, unless otherwise noted, were obtained from Sigma-Aldrich (St. Louis, MO).

### Cell culture and transfection

Human embryonic kidney (HEK) 293 cells were cultured in Dulbecco’s modified Eagle’s medium (DMEM) supplemented with 10% fetal bovine serum, penicillin (100 U/ml), and streptomycin (100 mg/ml). Chinese hamster ovary (CHO)-K1-EA cells were cultured in Ham’s F12 media supplemented with 10% fetal bovine serum, penicillin (100 U/ml), streptomycin (100 mg/ml), hygromycin (300 μg/ml) and G418 (800 μg/ml). Cells were grown at 37°C in 5% CO_2_ and 90% humidity. HEK293 cells were seeded in 100- or 35-mm plates and transfected overnight using a 1:3 ratio [1 µg of DNA:3 ml of polyethyleneimine (PEI)] diluted to 1 ml in non-supplemented DMEM and added to the cells already in non-supplemented DMEM. Media were replaced with culture media the following day. Concentrations of DNA are indicated for each experiment type. The cells were routinely checked and found to be negative for mycoplasma infection.

### Radioligand binding assays

Radioligand competition binding assays were conducted with slight modifications as previously described by our laboratory^51–53^. CHO-K1 cells stably expressing the human D1R, D2R, D3R, D4R, or D5R, or HEK293 cells transiently expressing the D2R, or D2R mutants, were dissociated from plates using Cell stripper (CHO cells; Corning, Inc., Corning, NY) or Earle’s balanced salt solution (EBSS) lacking calcium and magnesium. Intact cells were collected by centrifugation at 1000g for 10 min. Cells were resuspended and lysed using hypotonic lysis buffer (5 mM Tris-HCl and 5 mM MgCl_2_ at pH 7.4) at 4°C. Cell lysates were pelleted by centrifugation at 30,000g for 30 minutes and resuspended in EBSS + CaCl_2_ at pH 7.4. In some experiments, as indicated, the final membrane pellet was resuspended in Tris buffer, pH 7.4 at 25°C, either with or without 140 mM NaCl. Membrane homogenates (100 μl, containing ∼10 to 20 μg of protein, quantified by the Bradford assay) were incubated for 90 min at room temperature with 0.2-0.3 nM [^3^H]-methylspiperone (for D2R, D3R and D4R) or 0.5 nM [^3^H]-SCH23390 (for D1R and D5R) and the indicated concentrations of drug in a final reaction volume of 250 μl. Nonspecific binding was determined in the presence of 4 μM (+)-butaclamol. Bound radioligand was separated from free by filtration through a PerkinElmer UniFilter-96 Harvester (PerkinElmer, Waltham, MA), and washed three times with ice-cold assay buffer filling the well each time. After drying, 50 μl of liquid scintillation cocktail (MicroScint PS; PerkinElmer) was added to each well, and the plates were sealed and radioactivity was quantified using a PerkinElmer Topcount NXT.

### DiscoverX β-arrestin recruitment assay

Assays were conducted with minor modifications as previously published by our laboratory^51, 52^ using the DiscoverX PathHunter technology (DiscoverX, Inc., Fremont, CA). Briefly, CHO-K1-EA cells stably expressing β-arrestin fused to an N-terminal deletion mutant of β-galactosidase and human D2R fused to a complementing N-terminal fragment of β-galactosidase (DiscoverX, Inc.), were maintained in Ham’s F12 media supplemented with 10% fetal bovine serum, 100 U/ml penicillin, 100 μg/ml streptomycin, 800 μg/ml G418 and 300 μg/ml hygromycin at 37°C, 5% CO_2_ and 90% humidity. Cells stably expressing the D2R were seeded at a density of 2625 cells/well in 7.5 μl/well in 384-well clear-bottomed plates. After 16-24 hr of incubation, cells were treated with multiple concentrations of compound in PBS containing 0.2 mM sodium metabisulfite and incubated at 37°C for 90 min. Tropix Gal-Screen buffer and substrate (Applied Biosystems, Bedford, MA) were added to cells according to the manufacturer’s recommendations and incubated for 30-45 min at room temperature. Luminescence was measured on a Hamamatsu FDSS μCell reader. Data were collected as relative luminescence units and subsequently normalized to the control compound as indicated in each figure. The Hill coefficients of the concentration-response curves did not differ from unity with the data fitting to a single site model.

### cAMP accumulation assay

D2R-mediated inhibition of forskolin-stimulated cAMP production was measured by using the TR-FRET–based LANCE cAMP assay (PerkinElmer, Inc., Waltham, MA). CHO-K1 cells stably expressing the D2R were plated in Hank’s balanced salt solution (with CaCl_2_ and MgCl_2_) with 5 mM HEPES buffer and 0.2 mM sodium metabisulfite at a density of 1 × 10^6^ cells/ml in 5 μl per well in 384-well white-bottom plates. Compounds and forskolin were made in the same buffer. Immediately after plating, cells were treated with 2.5 μl of varying concentrations of compound and 2.5 μl of forskolin (10 μM final concentration) and incubated for 30 min at room temperature. Eu-cAMP tracer (5 μl) and ULight-anti-cAMP (5 μl) solutions were added to each well according to the manufacturer’s protocol, and cells were incubated in the dark for 2 hr at room temperature. Plates were read on a PHERAstar plate reader (BMG LABTECH, Cary, NC) with excitation at 337 nm and emission at 620 and 665 nm. Data were obtained as the ratio between A (excitation at 337 nm/emission at 665 nm) and B (excitation at 337 nm/emission at 620 nm). Values were normalized to a percentage of the control TR-FRET signal seen with a maximum concentration of dopamine. The Hill coefficients of the concentration-response curves did not significantly differ from unity with the data fitting to a single site model.

### BRET Assays

HEK293 cells were seeded at a density of 4 x 10^6^ cells per 100-mm dish and incubated overnight. The next day, cells were transiently transfected with the indicated constructs using PEI (DNA:PEI, 1:3 ratio). Go BRET assays were transfected with 0.5 μg G*_α_*_o1_-RLuc8, along with 5 μg G_γ2_-mVenus, 4 μg G_β1_, and 5 μg of the corresponding untagged D2R. β-arrestin assays were transiently transfected with 1 μg D2R-Rluc8 and 5 μg β-arrestin2-mVenus. Forty-eight hr after transfection, cells were harvested, washed, and resuspended in Dulbecco’s phosphate-buffered saline (DPBS) containing 0.2 mM sodium metabisulfite and 5.5 mM glucose. Cells were then plated in 96-well, white, solid-bottomed plates (Greiner Bio-One) at 100,000 cells/well and incubated in the dark for 45 min. Afterwards, curve shift assays were treated with indicated compounds for 90 min and experiments assessing inverse agonist activity of antagonists were treated with indicated compound for 30 min. Following compound incubation, cells were treated with 5 μM coelenterazine-H (NanoLight^TM^ Technology) for 5 min and analyzed with a PHERAstar plate reader (BMG LABTECH, Cary, NC). BRET signals were determined by calculating the ratio of the light emitted by mVenus (535/30 nm) over that emitted by Rluc8 (475/30 nm). Net BRET values were obtained by subtracting the background ratio from vehicle-treated wells. Experiments involving concentration-response curves were fit to non-linear regression analysis to determine the EC_50_ or IC_50_ values.

### Psychoactive Drug Screening Program (PDSP) panel

ML321 was screened using the National Institute of Mental Health Psychoactive Drug Screening Program (PDSP) directed by Dr. Bryan L. Roth (University of North Carolina, Chapel Hill, NC) ^32^. A complete list of the targets in this assay is presented in **Table S1**. For experimental details including radioligands used and associated Kd values for each individual target, please refer to the PDSP website http://pdsp.med.unc.edu/. Screening was performed using 10 µM of the test compound and >50% inhibition of radioligand binding is considered significant. Assays were performed in triplicate.

### DiscoverX gpcrMAX^TM^ GPCR Panel

As part of our assessment of the receptor selectivity of ML321, we used the DiscoverX gpcrMAX GPCR panel, which measures β-arrestin recruitment to different GPCRs (see DiscoverX β-arrestin recruitment assay methods above). This study was conducted by DiscoverX, Inc. (Fremont, CA). β-arrestin recruitment to each GPCR in the panel was stimulated by an agonist for that specific GPCR in the presence of 10 μM ML321 for antagonist mode assays, or 10 μM ML321 alone for agonist mode assays. Assay results, run in duplicate, are presented as the mean percent inhibition or stimulation of the indicated GPCR for each compound tested. Responses that deviate >20% from baseline are considered significant. For a full description of the DiscoverX gpcrMAX GPCR panel, see http://www.DiscoverX.com.

### Molecular modeling and molecular dynamics simulations

We docked the ML321 molecule into our previously established D2R model ^38, 54^ with the induced-fit docking protocol^55^ implemented in Schrödinger suite (Schrödinger, LLC: New York, NY). From the initial docking results, we chose four slightly different ML321 poses that have its dibenzothiazepine moiety bound in the orthosteric binding site (OBS) with the amide nitrogen forming a hydrogen bond interaction with the sidechain carboxyl group of Asp114^3.32^ and the thiophene moiety largely oriented towards the secondary binding pocket formed by TMs 2, 3, and 7. Each of the selected D2R-ML321 complex models were then immersed in the explicit water and 1-palmitoyl-2-oleoylphosphatidylcholine (POPC) lipid bilayer environment. The system charges were neutralized and 150 mM NaCl was added. The total system size was ∼75,000 atoms. We used the CHARMM36 force field^56, 57^ together with TIP3P water model. The ML321 parameters were obtained through the GAAMP server^58^ with the initial force field based on CGenFF assigned by ParamChem^59^. The MD simulations were carried out with Desmond (version 3.8; D. E. Shaw Research, New York, NY) as described previously^38, 54, 60^. Each system was first minimized for 6000 steps with restraints on all the heavy atoms of the ligand and protein, then for 6000 steps without any restraints. The following equilibration was performed with restraints on the heavy atoms of ligand and protein backbone in two stages, first in a canonical (NVT) ensemble for 60 ps, then in an isothermal–isobaric (NPT) ensemble for 600 ps. The production runs were carried out in an NPT ensemble with all atoms unrestrained. The NPT ensemble at 310 K and 1 atm was maintained by Langevin dynamics. For each of the four MD systems, we collected at least one 360 ns trajectory, and they all converged to the pose presented in the text. For a representative system, we collected two 480 ns trajectories, and used one of them to generate a video (**Supplemental Video**).

### Measurement of D2R binding kinetics using a time-resolved fluorescence resonance energy transfer assay (TR-FRET)

The association and dissociation rates of ligand binding to the D2R using a TR-FRET assay were determined as previously described^20^. The PPHT ((±)-2-(n-phenethyl-n-propyl)amino-5-hydroxytetralin hydrochloride;1-Naphthalenol, 5,6,7,8-tetrahydro-6-[(2-phenylethyl)propylamino]) derivative labeled with a red fluorescent probe (PPHT-red) was obtained from Cisbio Bioassays (Bagnolssur-Cèze, France). Briefly, CHO cells were stably transfected with a SNAP-tagged human D2SR (short isoform) and maintained in Dulbecco’s modified Eagle’s medium (DMEM) containing 2 mM glutamine (Sigma-Aldrich, Poole, UK) and supplemented with 10% fetal calf serum (Life Technologies, Paisley UK). Cells were incubated with Tag-lite labeling medium (LABMED) containing 100 nM of SNAP-Lumi4-Tb for one hour at 37°C followed by washing, detachment, disruption, and preparation of membranes, which were frozen at −80°C until use. Protein concentration was determined using the bicinchoninic acid assay kit (Sigma-Aldrich). Prior to their use, the frozen membranes were thawed and suspended in the assay buffer at a protein concentration of 0.2 mg/ml.

All fluorescent binding experiments using PPHT-red were conducted in white 384-well Optiplate plates in assay binding buffer, 20 mM HEPES: 138 mM NaCl, 6 mM MgCl2, 1 mM EGTA, and 1 mM EDTA and 0.02% pluronic acid (pH 7.4), 100 μM GppNHp, and 0.1% ascorbic acid. GppNHp was included to remove the G protein-coupled population of receptors that can result in two distinct populations of binding sites in membrane preparations, since the Motulsky–Mahan model^61^ is only appropriate for ligands competing at a single site. This was necessary because PPHT-red is an agonist. In all cases, nonspecific binding was determined in the presence of 10 μM haloperidol.

To accurately determine association rate (*k*_on_) and dissociation rate (*k*_off_) values, the observed rate of association (*k*_ob_) was calculated using at least four different concentrations of PPHT-red. The appropriate concentration of PPHT-red was incubated with human SNAP-D2R_S_ CHO cell membranes (2 μg per well) in assay binding buffer (final assay volume, 40 μl). The degree of PPHT-red bound to the receptor was assessed at multiple time points by HTRF detection to allow construction of association kinetic curves. The resulting data were globally fitted to the following equation (Equation 1) to derive a single best-fit estimate for *k*_on_ and *k*_off_:

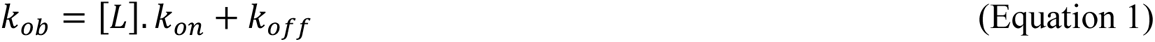

From these analyses we obtained the following parameters for the binding of PPHT-red to the hD2SR: *k*_off_ = 0.23 ± 0.01 min^-1^, *k*_on_ = 2.4 ± 0.31 x10^7^ M^-1^ min^-1^, *K*_d_ (*k*_off_ / *k*_on_) of 11.5 ± 0.16 nM, which are in good agreement with our previously published values for these parameters. Data are the mean ± SEM of 11 independent experiments performed in triplicate.

To determine ligand association and dissociation rates, we used a competition kinetic binding assay involving the simultaneous addition of both fluorescent ligand and competitor to the receptor preparation, so that at *t* = 0 all receptors are unoccupied. PPHT-red at 12.5 nM (a concentration which avoids ligand depletion in this assay volume) was added simultaneously with the unlabeled compound (at *t* = 0) to CHO cell membranes containing the human D2R_S_ in 40 μl of assay buffer. The degree of PPHT-red bound to the D2R_S_ was assessed at multiple time points by HTRF detection on a Pherastar FS (BMG Labtech, Offenburg, Germany) using standard HTRF settings. The terbium donor was always excited with three laser flashes at a wavelength of 337 nm. A kinetic TR-FRET signal was collected at 20-sec intervals both at 665 and 620 nm. HTRF ratios were obtained by dividing the acceptor signal (665 nm) by the donor signal (620 nm) and multiplying this value by 10,000. Nonspecific binding was defined as the amount of HTRF signal detected in the presence of haloperidol (10 μM) and was subtracted from each time point. Each time point was conducted on the same 384-well plate incubated at 37°C with orbital mixing. Multiple concentrations of unlabeled competitor were tested for determination of rate parameters. These data were globally fitted using the following equation (Equation 2) described by Motulsky & Mahan^61^ to simultaneously calculate *k*_on_ and *k*_off_ for unlabeled clozapine and ML321:

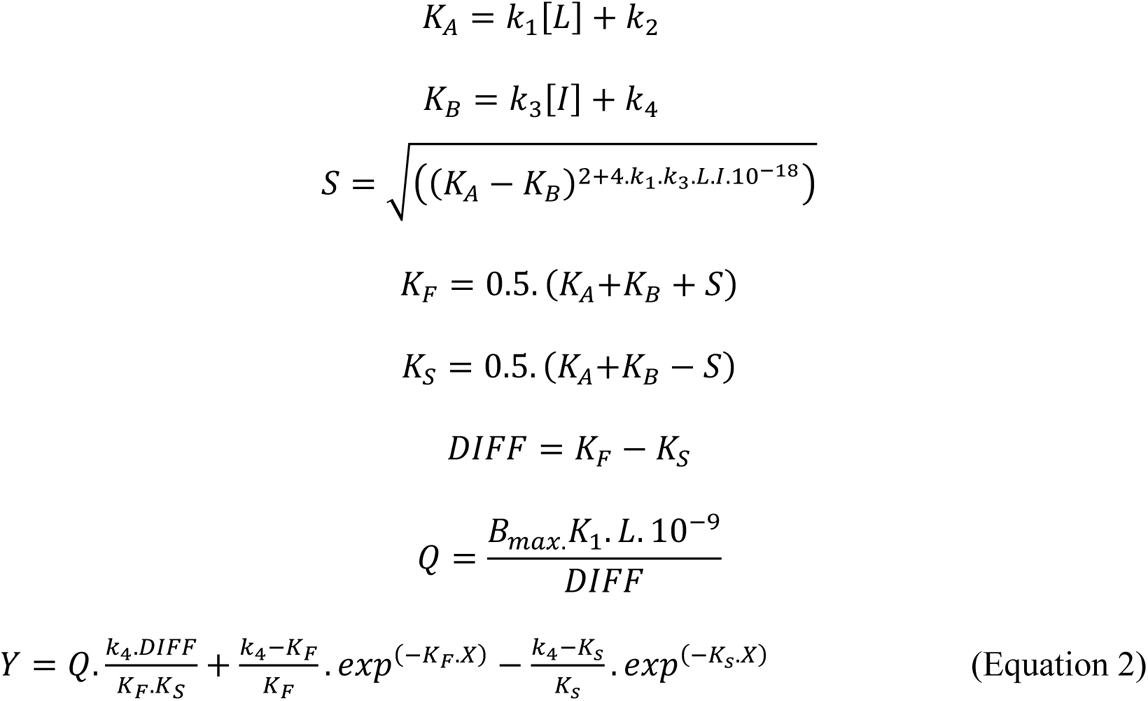

Where: X = time (min), Y = specific binding (HTRF ratio 665 nm/620 nm x 10,000), k_1_ = *k*_on_ PPHT-red (M^-1^ min^-1^), *k*_2_ = *k*_off_ PPHT-red (min^-1^), L = concentration of PPHT-red used (nM), Bmax = total binding (HTRF ratio 665 nm/620 nm x 10,000), I = concentration of unlabeled antagonist (nM). Fixing the above parameters allowed the following to be calculated: *k*_3_ = association rate of unlabeled ligand (M−1 min−1), *k*_4_ = dissociation rate of unlabeled ligand (min−1). The dissociation half-life of the various compounds was obtained using the following equation (Equation 3):

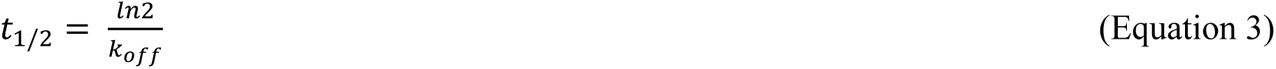

### PET data acquisition

MicroPET imaging studies were conducted as previously described^42^ using a Focus 220 microPET scanner (Siemens Medical Systems, Knoxville, TN, USA). Male rhesus monkeys (8–12 kg) were initially anesthetized with ketamine (10–15 mg/kg) and injected with glycopyrrolate (0.013–0.017 mg/kg) to reduce saliva secretions. The PET tracer was administered (i.v.) ∼90 min after ketamine injection. Subjects were intubated and placed on the scanner bed with a circulating warm-water blanket. Anesthesia was maintained with isoflurane (1.0–1.75% in 1.5 L/min oxygen flow). Respiration rate, pulse, oxygen saturation, body temperature, and inspired/exhaled gases were monitored throughout the study. Radiotracers and fluids were administered using a catheter placed percutaneously in a peripheral vein. In each microPET scanning session, the head was positioned supine with the brain in the center of the field of view. A 10-min transmission scan was performed to check positioning; once confirmed, a 45-min transmission scan was obtained for attenuation correction.

Subsequently, a 100-min baseline dynamic emission scan was acquired after administration of ∼10 mCi of [^11^C]SV-III-130 via the venous catheter. Displacement studies were also conducted in animals by administering ML321 at 20-min post-tracer injection. Acquired list mode data were transformed into a 3D set of sinograms and binned to the following time-frames: 3 x 1 min, 4 x 2 min, 3 x 3 min, and 20 x 5 min. Sinogram data were corrected for attenuation and scatter. Maximum *a posteriori* (MAP) reconstructions were performed with 18 iterations and a beta value of 0.004, resulting in a final 256 x 256 x 95 matrix. The voxel dimensions of the reconstructed images were 0.95 x 0.95 x 0.80 mm. A 1.5 mm Gaussian filter was applied to smooth each MAP reconstructed image. These images were then co-registered with MRI images to identify the brain regions of interest to obtain time-activity curves. All studies were conducted in accordance with the Guide for the Care and Use of Laboratory Animals^62^, and experimental procedures were approved by the University of Pennsylvania Committee on the Use and Care of Animals.

### Behavioral Experimentation

#### Subjects

The hyperthermia and yawning experiments were conducted with male Sprague-Dawley rats (Harlan Laboratories; Indianapolis, IN) weighing at least 250 g. Rats were housed 3 per cage with *ad libitum* access to tap water and standard laboratory chow in a temperature- and humidity-controlled facility under a 12-hr light-dark cycle (lights on at 0700 hr). Animals were allowed to acclimate to the facility for at least 5 days and were acclimated to handling for 2-3 days before testing. All rat experiments were conducted during the light phase of the cycle with naïve animals (i.e., each animal experienced only one experimental condition, and all dose comparisons were made between subjects). Adult male and female C57BL/6J mice (Jackson Laboratories, Bar Harbor, ME) were tested in the open field, prepulse inhibition (PPI), and catalepsy studies and had been acclimated to the facility for at least 2 weeks before testing. The naïve mice were housed 3-5 mice/cage on a 14:10 hr light/dark cycle (lights on 0700 hr) in a humidity- and temperature-controlled room with food and water provided *ad libitum*. The mouse behavioral experiments were conducted during the light cycle (0900-1600 hr). All studies were performed in accordance with the Guide for the Care and Use of Laboratory Animals^62^, and all experimental procedures were approved by the University of Michigan Committee on the Use and Care of Animals (rats) or the Duke University Institutional Animal Care and Use Committee (mice).

#### Drugs

For the rat studies, sumanirole was obtained from Tocris Bioscience (Bristol, UK) or Haoyuan Chemexpress (Shanghai, China). Pramipexole was obtained from APAC Pharmaceutical (Columbia, MD). ML321 was dissolved in a vehicle of 10% (v/v) *N*-methyl-2-pyrrolidone (NMP) + 20% (v/v) PEG400 + 70% (v/v) 25% (w/v) β-cyclodextrin in water with sonication until clear. All ML321 solutions were prepared directly before use. Pramipexole and sumanirole were dissolved in physiological saline. For the rat studies, all drugs were injected (s.c.) in a volume of 1.0 ml/kg except for 10.0 mg/kg ML321, which was injected in a volume of 3.125 ml/kg. For the mouse studies, amphetamine (AMPH), phencyclidine (PCP), and haloperidol (HAL) were purchased from Sigma-Aldrich. The ML321 was dissolved in a small amount of *N,N*-dimethylacetamide (DMA) with previously heated Tween-80 (Sigma-Aldrich) and vortexed until the solution was solubilized. Sterile-filtered water (Corning Meditech, Manassas, VA) was added to the mixture to provide a final concentration of 5% DMA with 5% Tween-80. The ML321 was administered (i.p.) in a volume of 10 ml/kg; all other drugs were given (i.p.) in a volume of 5 ml/kg.

#### Equipment

For the rat studies, experiments were conducted in transparent plastic observation chambers (48 x 23 x 20 cm), which resembled the animals’ home-cages, except there was no food, water, or bedding in the observation chambers. For the observation of yawning, angled mirrors were placed behind the chambers to facilitate viewing the rat regardless of its position in the chamber. For the mouse experiments, motor activity was assessed in a 21 x 21 x 30 cm^3^ open field (Omnitech Inc., Columbus, OH), and PPI was tested in a San Diego Instruments apparatus (San Diego, CA) as described previously^63^. Catalepsy was evaluated in two separate formats; it was assessed with the horizontal bar (6 mm wooden pole, 30 cm long, and 3 cm high) as described^63^, and with the 45° inclined screen [the black polyester PetScreen was 16 mesh, 45 x 50 cm in size, and bordered with an aluminum frame (Phifer Inc., Tuscaloosa, Alabama)] assay as outlined^64^.

#### Observation of hypothermia

On test day, each rat was placed into an observation chamber and allowed to acclimate for 30 min. After acclimating, the first (i.e., pre-drug baseline) rectal temperature was taken. Temperature measurements were obtained using a digital veterinary thermometer (model Adtemp 422; American Diagnostic Corporation; Hauppauge, NY) with the rats manually restrained and the thermometer inserted 2-3 cm into the rectum. Temperature readings (°C) were obtained within ∼10 sec. Baseline temperatures were normal for the rat (mean ± SEM: 37.4 ± 0.1). Immediately after the first temperature measurement, rats were injected (s.c.) with vehicle or 1.0 mg, 3.2 mg, or 10.0 mg/kg of ML321 and returned to the observation chamber. Thirty min later, rats were injected (s.c.) with vehicle or 1.0 mg/kg sumanirole and returned to the observation chamber. The second (i.e., post-drug) rectal temperature was taken 30 min after vehicle/sumanirole administration. The change in each animal’s body temperature was calculated as the difference between the baseline (first) and post-drug (second) temperatures.

#### Observation of yawning

Each rat was placed into an observation chamber and allowed to acclimate for 30 minutes. After acclimating, rats were injected (s.c.) with vehicle or 3.2 mg/kg or 10.0 mg/kg of ML321 and returned to the observation chamber. Thirty min later, rats were injected (s.c) with vehicle or 0.1 mg/kg pramipexole, returned to the chamber, and observed for 60 minutes. The number of unique yawns, defined as a prolonged (∼1 sec) wide opening of the mouth followed by a rapid closure, was recorded by a trained observer.

#### Locomotor Activity

Motor activity in mice was assessed in a 21 x 21 x 30 cm^3^ open field (Omnitech Inc., Columbus, OH) as described^63^. The mice were housed in the test room 24 hr prior to testing. Animals were injected (i.p) with the vehicle or different doses of ML321 and immediately placed into the open field. After 30 min, the mice were removed and injected with the vehicle, 3 mg/kg AMPH, or 6 mg/kg PCP and returned immediately to the open field for an additional 90 min. Horizontal distance traveled (cm) was quantitated with Fusion software (Omnitech) and scored in 5-min bins across testing or as cumulative locomotor activity at baseline (0-30 min) or post-injection (31-120 min).

#### Prepulse Inhibition (PPI)

Mice were administered (i.p.) the vehicle or various doses of ML321 in their home cage. Ten min later, animals were given the vehicle, 3 mg/kg AMPH, or 6 mg/kg PCP and acclimated to a 65 dB white-noise background in the PPI apparatus (San Diego Instruments, San Diego, CA). Five min later, mice were presented with combinations of startle (120 dB), prepulse-pulse (4, 8, and 12 dB over the 65 dB background followed by 120 dB), and null trials over 25 min as described^63^. Activity was recorded during all trials. The data are presented as percent PPI = [1 − (prepulse trials/startle-only trials)] x 100, and as startle and null activities.

#### Catalepsy

Catalepsy was assessed using the horizontal bar test as described^63^. Baseline responses were recorded and separate groups of mice were injected (i.p.) with either vehicle or various doses of HAL or ML321. Mice were tested 60 min later for catalepsy. The latency for a mouse to remove its paws from the bar was recorded as an index of catalepsy with a maximum time of 60 sec. Catalepsy was evaluated also by the inclined screen test^64^. Mice were injected (i.p.) with vehicle or 10 mg/kg of either haloperidol (HAL) or ML321 (ML). Animals were returned to their home-cages and then tested 30 and 60 min later for catalepsy. Here, the mice were initially placed face-down on a horizontal wire-mesh screen that was inclined at a 45° angle and the latency to move its four paws, or one body length was recorded with a 300 sec cut-off.

### hERG Channel Assay

The study was performed by Cyprotex (https://www.cyprotex.com/). HEK293 cells expressing the hERG potassium channel were dispensed onto chips and hERG tail currents were measured by whole cell voltage clamping. A range of concentrations (up to 25 μM) of the test compound was added to the cells and a second recording of the hERG current was made. The percent change in hERG current was calculated and used to calculate an IC_50_ value. The experiment was performed on a Cytopatch Automated PatchClamp platform (Cytocentrics Inc.), which automatically performs electrophysiology measurements on cells on microchips. To confirm hERG sensitivity, 5.4 μM quinidine was applied to cells, blocking the hERG current by 99%.

### Cytotoxicity Screening Panel

The study was performed by Cyprotex (https://www.cyprotex.com/). HepG2 cells were plated on 384-well treated black-wall, clear-bottom polystyrene tissue culture plates. The cells were dosed with test compound using a range of concentrations. At the end of the incubation period, the cells were loaded with the relevant dye/antibody for each cell health marker. The plates were then scanned using an automated fluorescent cellular imager, ArrayScan® (Thermo Scientific Cellomics).

### Ames Mutagenicity Screening

The study was performed by Cyprotex (https://www.cyprotex.com/). Approximately ten million bacteria (*S. typhimurium*) were exposed in triplicate to test agent (six concentrations), a negative control (vehicle) and a positive control (2-aminoanthracene) for 90 minutes in medium containing a low concentration of histidine (sufficient for about 2 doublings.) The cultures were then diluted into indicator medium, lacking histidine, and dispensed into 48 wells of a 384-well plate (micro-plate format, MPF). The plate was then incubated for 48 hr at 37°C, and cells that had undergone a reversion will grow, resulting in a color change in wells. The number of wells showing growth were counted and compared to the vehicle control. An increase in the number of colonies of at least two-fold over baseline (mean + SD of the vehicle control) indicated a positive response.

### Data Analysis and Statistics

Data were analyzed using GraphPad Prism 9 (GraphPad Software, San Diego, CA). For the DiscoverX β-arrestin recruitment assay, the LANCE cAMP assay, and the BRET curve-shift assays, the data were fit using non-linear regression analysis. For curve-shift assays a Schild-type analysis was conducted using the equation y = log((A’/A)-1) where A’ was the EC_50_ of dopamine in the presence of ML321 and A was the EC_50_ of dopamine without ML321. The x-axis of the Schild graph was the molar concentration of ML321. Linear regression was performed on the resulting plot. Statistical significance was determined using one-way analysis of variance (ANOVA) with Dunnett’s *post-hoc* test when comparing to a control group. For the rat studies, the statistical analyses were performed with Prism 6.0 (GraphPad Software), and the results were presented graphically as the mean ± SEM. When ML321 was given as a pretreatment to 1 mg/kg sumanirole or 0.1 mg/kg pramipexole, mean temperature changes or mean total yawns, respectively, were analyzed by ML321 dose using one-way ANOVA for ML321 dose. Following a significant effect, pairwise comparisons to 0.0 mg/kg ML321 (vehicle) were made *post hoc* using Dunnett’s tests. When ML321 was given as a pretreatment to vehicle (0.0 mg/kg sumanirole or pramipexole), the data were presented for reference but were not formally analyzed due to the small number of animals (N=3/group) tested with 10 mg/kg ML321. To obtain a time-course of pramipexole-induced yawning, the 60-minute session was divided into 4 blocks, each encompassing 15 minutes, and the data were reanalyzed by ML321 dose and block using two-way ANOVA for ML321 dose and block. The mouse data were analyzed by SPSS programs (version 28) (IBM, Chicago, IL) and were presented as means and SEMs. The open field, PPI, and catalepsy data were analyzed by repeated measures ANOVA, while the null and startle activities from the PPI and the cumulative locomotor activities for the baseline and post-administration intervals were analyzed separately by one-way ANOVA. Terms significant in the parametric models were analyzed *post-hoc* by Bonferroni corrected pair-wise comparisons. For all statistics listed in this section, *post-hoc* comparisons were two-tailed and a *p*<0.05 was considered statistically significant.

## Supporting information

Supplemental MD simulation

## AUTHORSHIP CONTRIBUTIONS

*Participated in research design*: DRS, RBF, ANN, NMB, TBD, RMR, MM, KHL, JWB, JX, HDL, RHM, JHW, JRL, LS, WCW.

*Conducted experiments*: ANN, NMB, TBD, RMR, VMP, MM, KHL, JWB, JX, HDM, LS.

*Contributed new reagents or analytic tools*: ADG, JJM

*Performed data analysis*: RBF, ANN, NMB, TBD, RMR, VMP, MM, KHL, JWB, JX, HDL, JHW, JRL, LS, WCW.

*Wrote or contributed to the writing of the manuscript*: DRS, RBF, ANN, NMB, RMR, MM, KHL, JWB, JX, JRL, LS, WCW.

The manuscript was read and approved by all authors.

## Notes

The authors declare no competing financial interests.

Data availability: All data needed to evaluate the conclusions in the paper are present in the paper and/or the Supporting Information.

## ACKNOWLEDGMENTS

We wish to thank Ms. Jiechun Zhou at Duke University Medical Center for breeding and maintaining the C57BL/6J mice and Ms. Elizabeth Begej for helping with some of the behavioral testing of mice. We also thank Ms. Aylin Golaszewski, Ms. Emily Jackson, and Mr. Ryan Tarpey for their assistance with the observation of yawning and hypothermia in rats.

## ABBREVIATIONS

AC_50_: half-maximal activity concentration
AMPH: amphetamine
BRET: Bioluminescence resonance energy transfer
cAMP: cyclic adenosine monophosphate
CI: confidence interval
CNS: central nervous system
CRC: concentration-response curve
D1R: D1 dopamine receptor
D2R: D2 dopamine receptor
D3R: D3 dopamine receptor
D4R: D4 dopamine receptor
D5R: D5 dopamine receptor
EL2: extracellular loop 2
EPS: extrapyramidal side-effects
FDA: Food and Drug Administration
GPCR: G protein-coupled receptor
HAL: haloperidol
IND: investigational new drug
ip: intraperitoneal
MD: molecular dynamics
NIH: National Institutes of Health
OBS: orthosteric binding site
PCP: phencyclidine
PDSP: Psychoactive Drug Screening Program
PRAM: pramipexole
PPI: pre-pulse inhibition
SAR: structure-activity relationship
SBP: secondary binding pocket
SUM: sumanirole
TM: transmembrane
TR-FRET: time-resolved fluorescence resonance energy transfer
Veh: vehicle.

## For Table of Contents Only

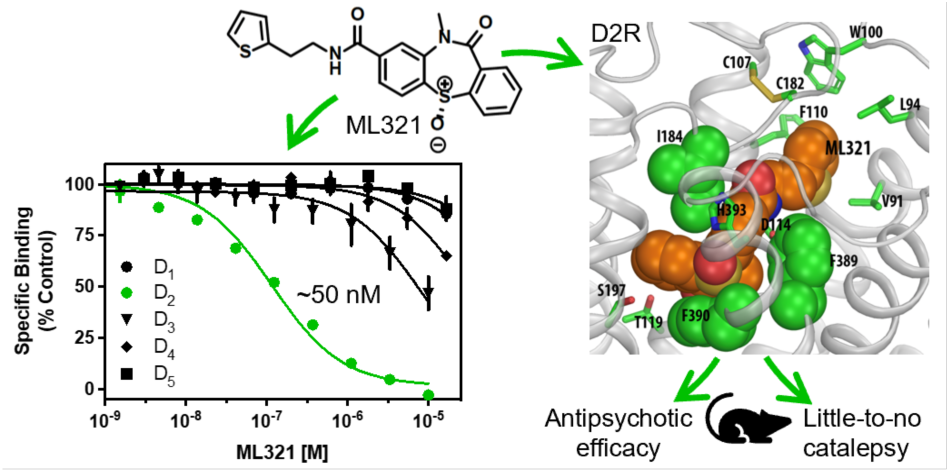

