## Supporting Information for "Identification and Characterization of ML321: a Novel and Highly Selective D*2* Dopamine Receptor Antagonist with Efficacy in Animal Models that Predict Atypical Antipsychotic Activity"

[a]Molecular Neuropharmacology Section, National Institute of Neurological Disorders and Stroke, Intramural Research Program, National Institutes of Health, 35 Convent Drive, MSC-3723, Bethesda, MD, 20892, United States. [b]Department of Psychiatry and Behavioral Sciences, Mouse Behavioral and Neuroendocrine Analysis Core Facility, Duke University Medical Center, 354 Sands Building, 303 Research Drive, Durham, NC, 27710, United States. [c]Computational Chemistry and Molecular Biophysics Section, Molecular Targets and Medications Discovery Branch, National Institute on Drug Abuse, Intramural Research Program, National Institutes of Health, 333 Cassell Drive, Baltimore, MD 21224, United States. [d]Department of Pharmacology, University of Michigan Medical School, 1150 W. Medical Center Dr., Ann Arbor, MI 48109, United States. [e]Division of Radiological Sciences, Department of Radiology, Mallinckrodt Institute of Radiology, Washington University School of Medicine, St. Louis, MO, USA. [f]Drug Discovery Biology, Monash Institute of Pharmaceutical Sciences, Monash University, 399 Royal Parade, Parkville, VIC 3052, Australia. [g]Division of Pre-Clinical Innovation, National Center for

Advancing Translational Sciences, National Institutes of Health, 9800 Medical Center Drive, Rockville, MD 20850, United States. [h]Department of Radiology, Perelman School of Medicine, University of Pennsylvania, 3400 Civic Center Boulevard, Philadelphia, PA 19104, United States. [i]Centre of Membrane Proteins and Receptors, Universities of Birmingham and Nottingham, Nottingham, NG7 2UH, United Kingdom. [j]Departments of Neurobiology and Cell Biology, Duke University Medical Center, 354 Sands Building, 303 Research Drive, Durham, NC, 27710, United States.

*Corresponding author:  David R. Sibley, Molecular Neuropharmacology Section, National Institute of Neurological Disorders and Stroke, Intramural Research Program, National Institutes of Health, 35 Convent Drive, MSC-3723, Bethesda, MD, 20892, United States..

**TABLE OF CONTENTS**

**Figure S1.** Functional profiling of ML321 against an array of 168 known GPCRs……….….S-4

**Figure S2.** ML321 is a competitive antagonist of D2R-mediated β-arrestin recruitment and Go protein activation…………………………………………………………………….…S-6

**Figure S3.** ML321 exhibits inverse agonist activity at the D2R in a concentration-dependent manner……………………………………………….…………………………………….S-7

**Figure S4.** Null and startle activities in the prepulse inhibition experiment shown in **Figure 10**…………………………………………………………………………..…………S-8

**Figure S5.** Catalepsy assessed using the inclined screen test……………………….........S-9

**Table S1.** Results from the Psychoactive Drug Screening Program (PDSP) assay…………. S-10

**Table S2.** Functional profiling of ML321 using the DiscoverX gpcrMAX$^{TM}$ assay panel as described in **Figure 2**……………………………………………………………………S-11

**Table S3.** Impact of D2R point mutations on ML321 antagonist potencies as assessed using dopamine-stimulated Go BRET activation assays……………………………………….S-12

**Table S4.** Impact of D2R point mutations on ML321 binding affinities as assessed using [$^3$H]-methylspiperone competition binding assays………………………………………………S-13

**Supporting Figures**

**Figure S1.** Functional profiling of ML321 against an array of 168 known GPCRs. The data are from **Figures 2A** and **B**, representing the panel screens with ML321 in antagonist and agonist modes, respectively. GPCRs of interest include $D2R_S$ (DRD2S), $D2R_L$ (DRD2L), D3R (DRD3), 5-$HT_{2A}$ serotonin (HTR2A), 5-$HT_{2C}$ serotonin (HTR2C), BLT1 (leukotriene $B_4$) (LTB4R), sphingosine-1-phosphate 4 (S1PR4), $\alpha_{2C}$-adrenergic (ADRA2C), and CB2 cannabinoid (CB2).

*Antagonist mode:*

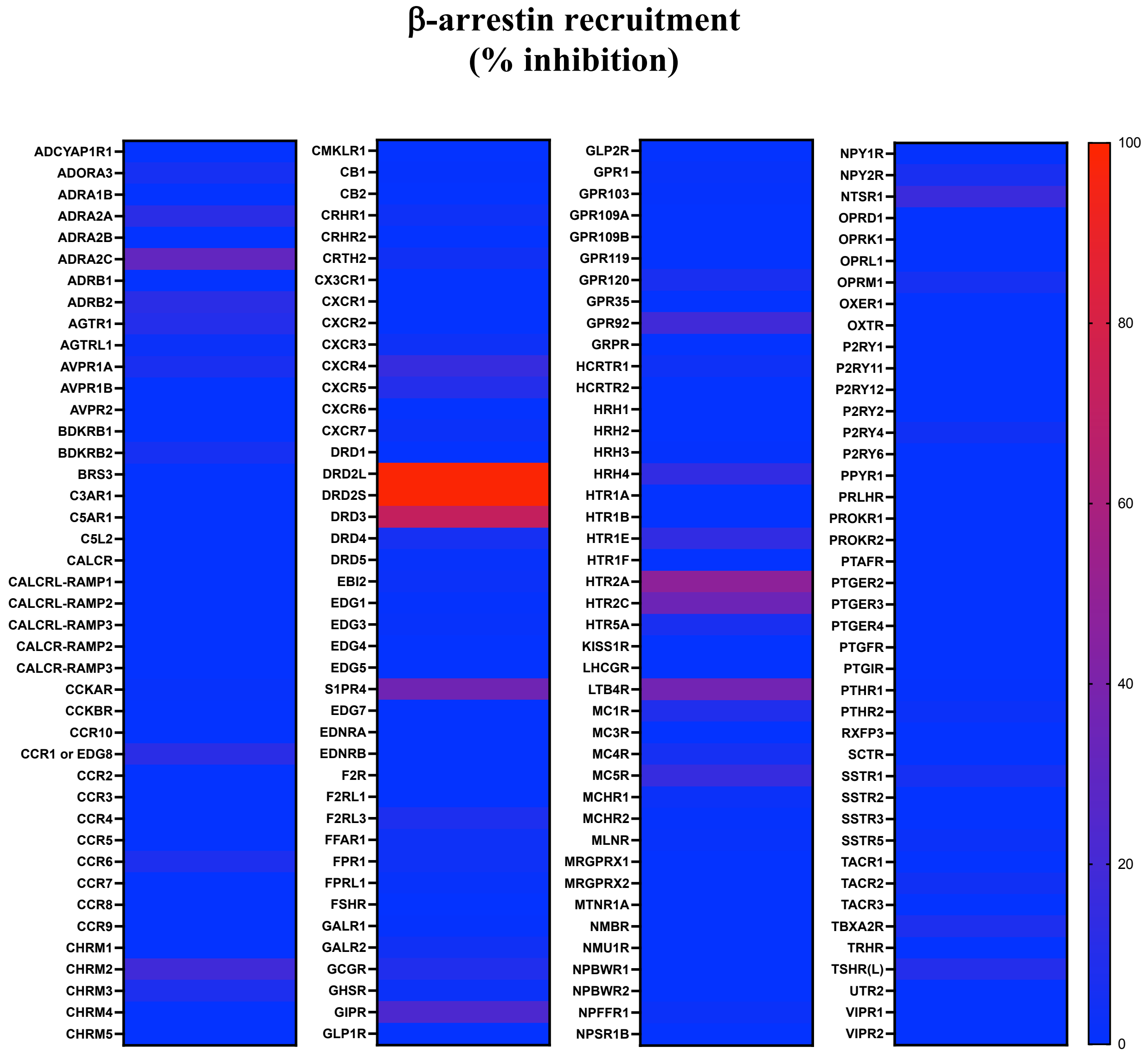

*Agonist mode:*

**β-arrestin recruitment**
**(% stimulation)**

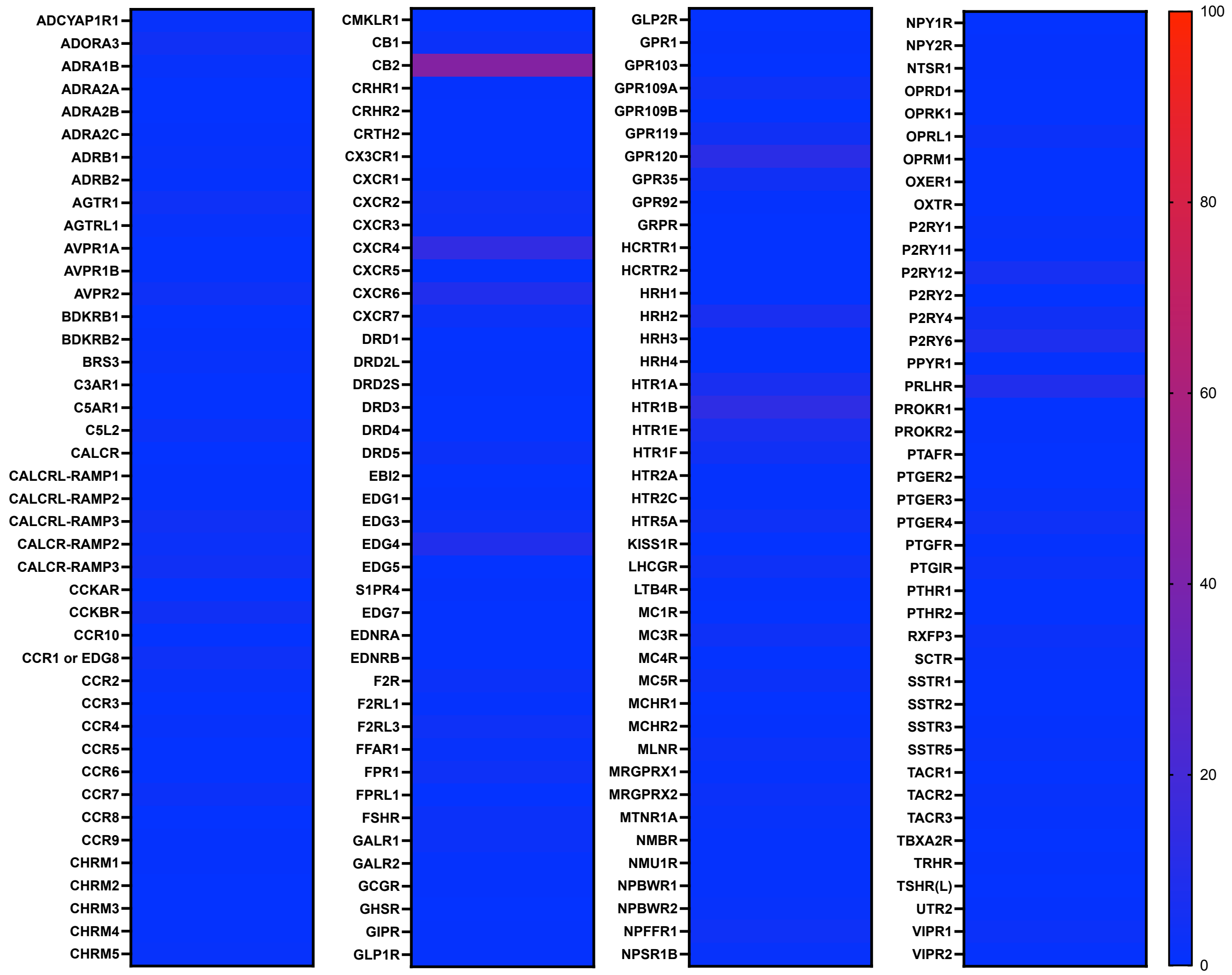

**Figure S2.** ML321 is a competitive antagonist of D2R-mediated β-arrestin recruitment and Go protein activation. Curve-shift assays were conducted by stimulating the receptor with the indicated concentrations of dopamine with or without various concentrations of ML321. Data are expressed as Δ BRET from baseline, which represents the constitutive BRET signal seen in the absence of any drug treatment. Insets represent Schild regression analyses of the data that were used to calculate mean $K_B$ values [95% C.I.]. (**A**) Dopamine-stimulated β-arrestin recruitment was assessed using a BRET-based assay as described in *Methods*. A representative experiment performed in triplicate is shown. ML321 $K_B$ = 239 nM [81-397] (N = 3). (**B**) D2R-mediated Go activation was measured using a BRET-based assay as described in *Methods*. A representative experiment performed in triplicate is shown. ML321 $K_B$ = 143 nM [10.1-441] (N = 3).

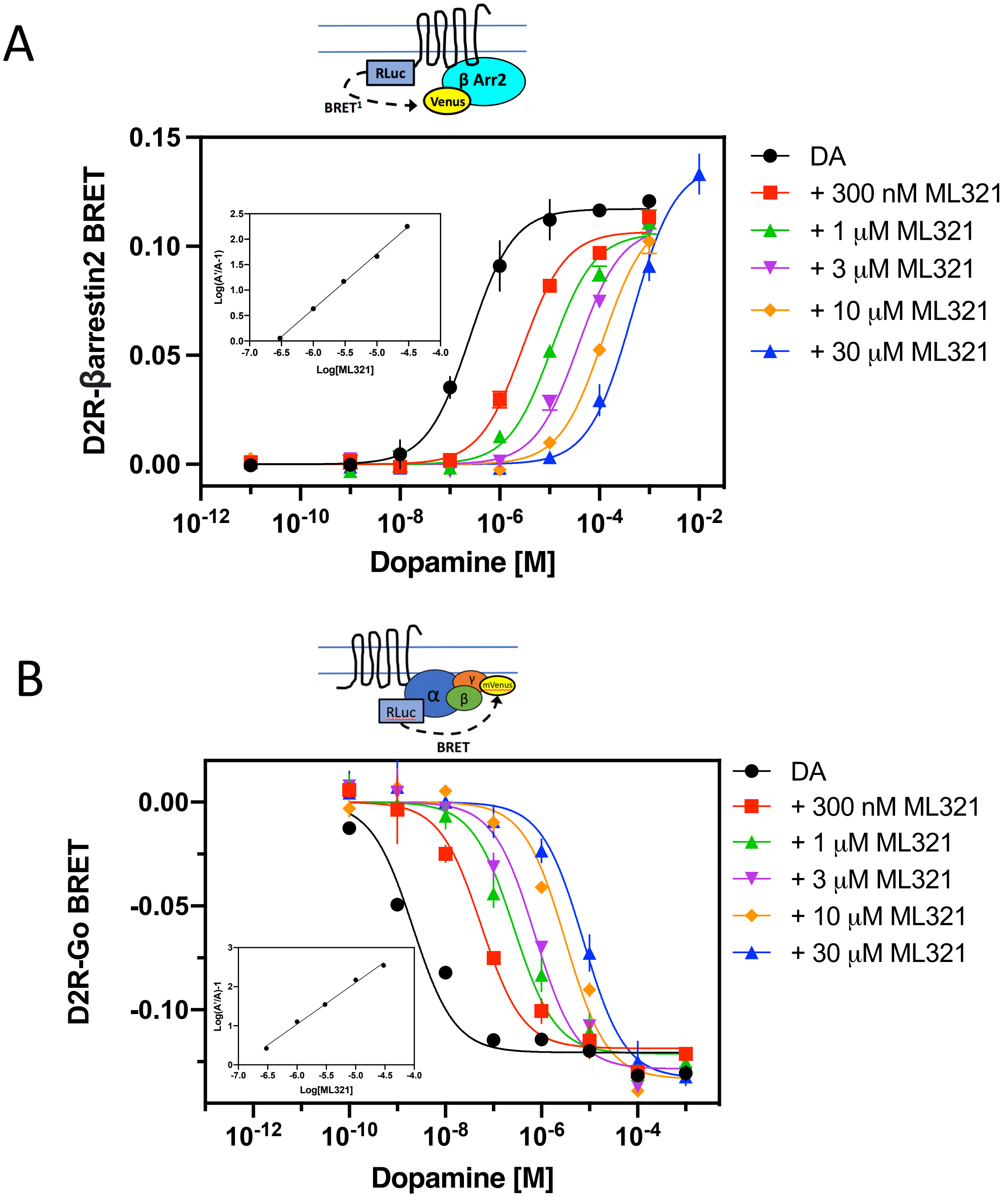

**Figure S3.** ML321 exhibits inverse agonist activity at the D2R in a concentration-dependent manner. Go BRET assays were performed as described in **Figure 4** (see legend). Data represent the mean ± SEM of five independent experiments performed in triplicate that are expressed as Δ BRET from baseline (zero), which represents the BRET signal seen in the absence of drug treatment. The mean ML321 $EC_{50}$ value [95% C.I.] = 275 nM [32 nM - 6 μM] (N = 5).

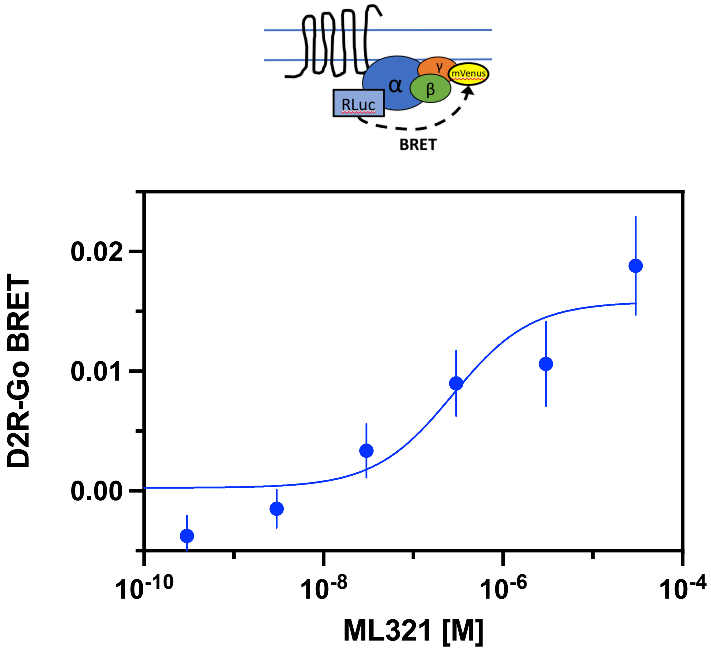

**Figure S4.** Null and startle activities in the prepulse inhibition experiment shown in **Figure 10**. The study was performed as described in the legend for **Figure 10** and in the *Methods*. (**A**) For null activity with amphetamine, a one-way ANOVA found treatment to be significant [$F(6,61)=8.165$, $p<0.001$]. **(B)** Similarly, for startle activity with amphetamine, the one-way ANOVA determined the treatment effect was significant [$F(6,61)=5.092$, $p<0.001$]. (**C**) In the phencyclidine experiment, a one-way ANOVA for null activity detected significant treatment effects [$F(6,61)=7.374$, $p<0.001$]. (**D**) For startle activities in the phencyclidine study, a one-way ANOVA identified the treatment effect to be significant [$F(6,61)=3.011$, $p=0.012$]. The data are presented as means ± SEMs. N=9 mice for the Veh-Veh, Veh-AMPH, and Veh-PCP groups; N=10 mice for all other groups. $^{\dagger}p<0.05$, for the AMPH startle study (startle activity) 5 mg/kg ML321(ML)-Veh *vs.* the Veh-Veh group; $^{*}p<0.05$, Veh-AMPH or Veh-PCP *vs.* the indicted groups; $^{\#}p<0.05$, 0.25 mg/kg ML-AMPH or ML-PCP *vs.* the indicated groups; $^{\wedge}p<0.05$, 0.5 mg/kg ML-AMPH or ML-PCP *vs.* the indicated groups; $^{+}p<0.05$, 1 mg/kg ML-AMPH or ML-PCP *vs.* the indicated groups; $^{\ddagger}p<0.05$, for the PCP study (null activity) 5 mg/kg ML-PCP *vs.* the Veh-Veh and 5 mg/kg ML-Veh groups.

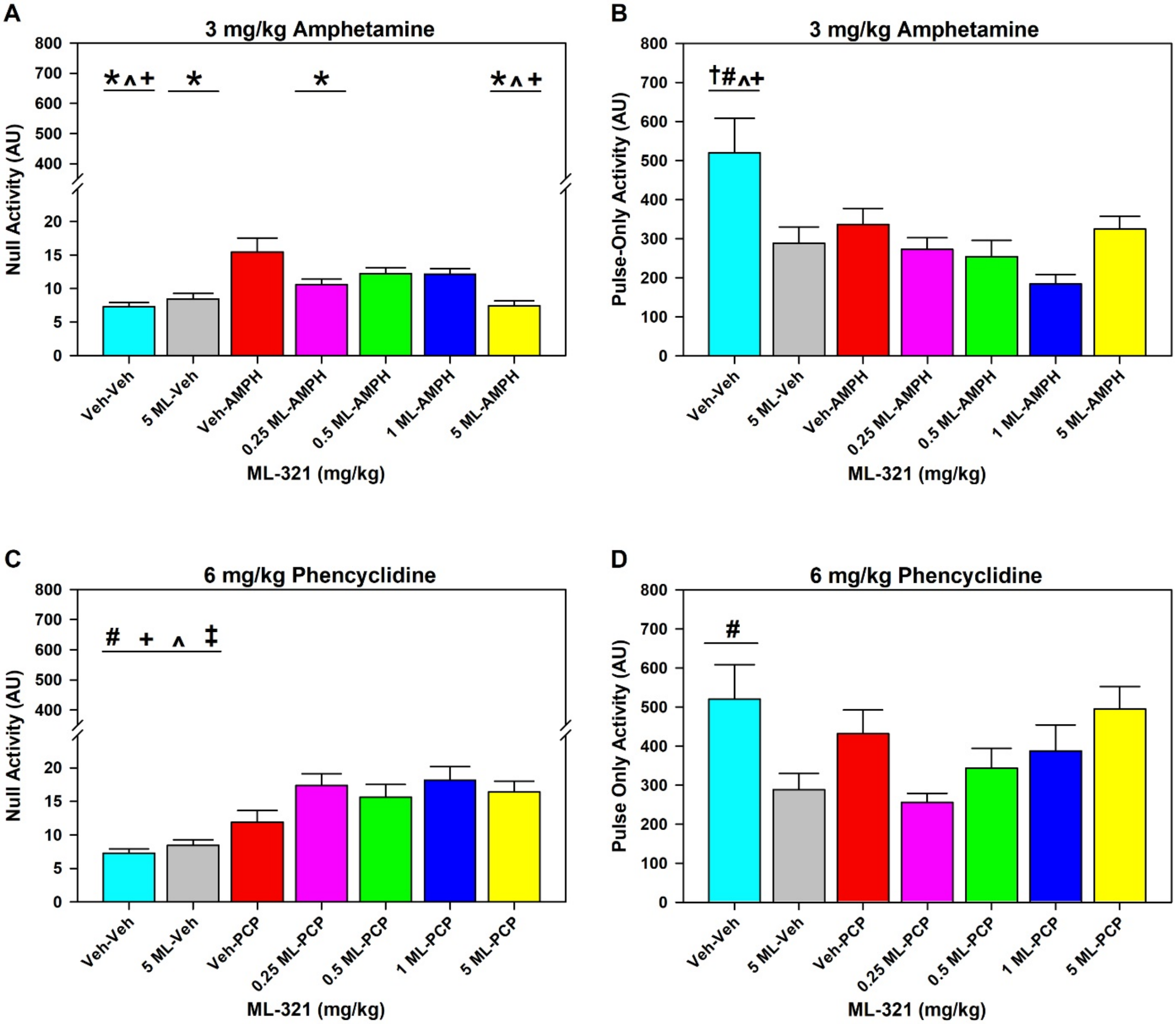

**Figure S5.** Catalepsy assessed using the inclined screen test. C57BL/6J mice were injected (i.p.) with vehicle or a 10 mg/kg dose of either haloperidol (HAL) or ML321 (ML). Mice were returned to their home cages and then tested 30 and 60 min later for catalepsy. In this test, the mice were initially placed face-down on a horizontal wire-mesh screen that was inclined at a 45° angle, and the latency to move the four paws (**A**) or one body length (**B**) was recorded with a 300 sec cut-off. The data are presented as means and SEMs. N=10 mice/group. A RMANOVA detected a significant treatment effect to move 4 paws [$F(1,18)=23.389$, $p<0.001$] and to move 1 body length [$F(1,18)=46.590$, $p<0.001$]. The data are presented as means ± SEMs. N=10 mice for all groups. *$p<0.05$, HAL *vs.* ML.

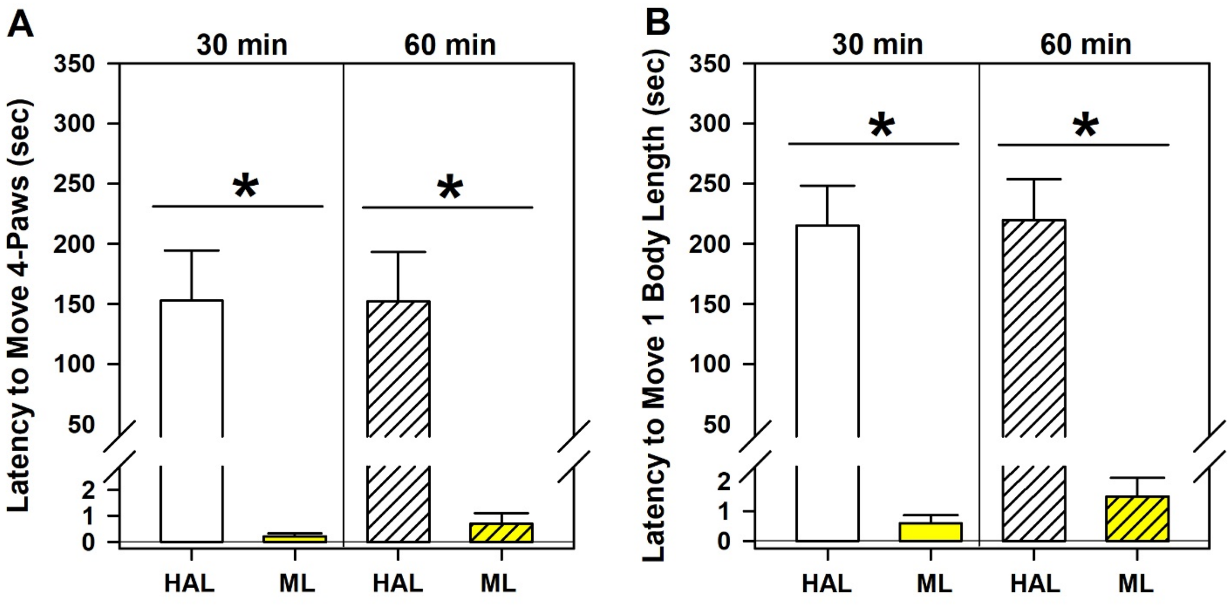

### Supporting Tables

**Table S1.** Results from the Psychoactive Drug Screening Program (PDSP) assay[a]

| **Target** | **ML321** (10 µM) **% Inhibition** | Target | **ML321** (10 µM) **% Inhibition** |
|---|---|---|---|
| 5HT1A | NA[b] | D2 | **92** |
| 5HT1B | NA | D3 | **59** |
| 5HT1D | NA | D4 | NA |
| 5HT1E | NA | D5 | NA |
| 5HT2A | NA | DAT | NA |
| 5HT2B | NA | DOR | NA |
| 5HT2C | **64** | $GABA_A$ | NA |
| 5HT3 | NA | H1 | NA |
| 5HT4 | NA | H2 | NA |
| 5HT5A | NA | H3 | NA |
| 5HT6 | NA | H4 | NA |
| 5HT7 | **53** | KOR | NA |
| Alpha1A | NA | M1 | NA |
| Alpha1B | NA | M2 | NA |
| Alpha1D | NA | M3 | NA |
| Alpha2A | NA | M4 | NA |
| Alpha2B | NA | M5 | NA |
| Alpha2C | NA | MOR | NA |
| Beta 1 | NA | NET | NA |
| Beta 2 | NA | PBR | NA |
| Beta 3 | NA | SERT | NA |
| BZP site | NA | Sigma 1 | NA |
| D1 | NA | Sigma 2 | NA |

[a] Radioligand binding assays were performed as described in the Methods. Data are from **Figure 1C** and represent the percent inhibition by ML321 of radioligand binding to each target.
[b] NA indicates inhibition of binding was <50% in the primary assay that was performed using a single (10 μM) concentration of ML321.

**Table S2.** Functional profiling of ML321 using the DiscoverX gpcrMAX[TM] assay panel as described in **Figure 2**. GPCRs of interest include $D2R_S$ (DRD2S), $D2R_L$ (DRD2L), D3R (DRD3), 5-$HT_{2A}$ serotonin (HTR2A), 5-$HT_{2C}$ serotonin (HTR2C), BLT1 (leukotriene $B_4$) (LTB4R), sphingosine-1-phosphate 4 (S1PR4), $\alpha_{2C}$-adrenergic (ADRA2C), and CB2 cannabinoid (CB2).

Panel key (GPCR list):

| | 1 | 2 | 3 | 4 | 5 | 6 | 7 | 8 | 9 | 10 | 11 | 12 | 13 | 14 | 15 | 16 | 17 | 18 | 19 | 20 | 21 |
|---|---|---|---|---|---|---|---|---|---|---|---|---|---|---|---|---|---|---|---|---|---|
| A | ADCYAP1R1 | AGTR1 | C3AR1 | CALCR-RAMP3 | CCR5 | CHRM4 | CX3CR1 | DRD1 | EDG3 | F2RL1 | GCGR | GPR10B | HRH1 | HTR2A | MC4R | NMBR | NTSR1 | P2RY11 | PROKR2 | PTHR2 | TACR2 |
| B | ADORA3 | AGTRL1 | C5AR1 | CCKAR | CCR6 | CHRM5 | CXCR1 | DRD2L | EDG4 | F2RL3 | GHSR | GPR119 | HRH2 | HTR2C | MC5R | NMU1R | OPRD1 | P2RY12 | PTAFR | RXFP3 | TACR3 |
| C | ADRA1B | AVPR1A | C5L2 | CCKBR | CCR7 | CMKLR1 | CXCR2 | DRD2S | EDG5 | FFAR1 | GIPR | GPR120 | HRH3 | HTR5A | MCHR1 | NPBWR1 | OPRK1 | P2RY2 | PTGER2 | SCTR | TBXA2R |
| D | ADRA2A | AVPR1B | CALCR | CCR10 | CCR8 | CB1 | CXCR3 | DRD3 | S1PR4 | FPR1 | GLP1R | GPR35 | HRH4 | KISS1R | MCHR2 | NPBWR2 | OPRL1 | P2RY4 | PTGER3 | SSTR1 | TRHR |
| E | ADRA2B | AVPR2 | CALCRL-RAMP1 | CCR1 or EDG8 | CCR9 | CB2 | CXCR4 | DRD4 | EDG7 | FPRL1 | GLP2R | GPR92 | HTR1A | LHCGR | MLNR | NPFFR1 | OPRM1 | P2RY6 | PTGER4 | SSTR2 | TSHR(L) |
| F | ADRA2C | BDKRB1 | CALCRL-RAMP2 | CCR2 | CHRM1 | CRHR1 | CXCR5 | DRD5 | EDNRA | FSHR | GPR1 | GRPR | HTR1B | LTB4R | MRGPR1 | NPSR1B | OXER1 | PPYR1 | PTGFR | SSTR3 | UTR2 |
| G | ADRB1 | BDKRB2 | CALCRL-RAMP3 | CCR3 | CHRM2 | CRHR2 | CXCR6 | EBI2 | EDNRB | GALR1 | GPR103 | HCRTR1 | HTR1E | MC1R | MRGPR2 | NPY1R | OXTR | PRLHR | PTGIR | SSTR5 | VIPR1 |
| H | ADRB2 | BRS3 | CALCR-RAMP2 | CCR4 | CHRM3 | CRTH2 | CXCR7 | EDG1 | F2R | GALR2 | GPR10A | HCRTR2 | HTR1F | MC3R | MTNR1A | NPY2R | P2RY1 | PROKR1 | PTHR1 | TACR1 | VIPR2 |

Antagonist Mode results from **Figure 2A**:

| | 1 | 2 | 3 | 4 | 5 | 6 | 7 | 8 | 9 | 10 | 11 | 12 | 13 | 14 | 15 | 16 | 17 | 18 | 19 | 20 | 21 |
|---|---|---|---|---|---|---|---|---|---|---|---|---|---|---|---|---|---|---|---|---|---|
| A | -2% | 10% | -10% | -7% | 0% | -3% | -2% | -5% | 2% | -2% | 9% | -11% | -9% | 48% | 6% | -14% | 17% | -2% | -9% | 3% | 5% |
| B | 6% | 3% | -1% | 2% | 8% | -14% | 0% | 98% | -7% | 8% | 3% | -3% | -13% | 35% | 16% | -5% | -3% | -14% | -4% | -4% | -2% |
| C | -1% | 7% | -6% | -3% | -5% | -4% | -24% | 98% | -10% | 4% | 23% | 7% | 1% | 7% | 3% | -9% | -7% | -7% | -3% | -11% | 8% |
| D | 12% | -6% | -14% | 0% | -12% | 1% | 4% | 72% | 36% | 4% | -7% | -8% | 14% | -3% | 1% | -2% | -9% | 5% | -8% | 6% | -18% |
| E | -16% | -8% | -6% | 12% | -3% | -7% | 16% | 6% | -4% | 2% | -6% | 19% | -1% | -9% | 2% | 3% | 6% | -8% | -2% | -6% | 10% |
| F | 31% | -9% | -4% | -6% | -1% | 4% | 10% | 2% | -2% | -25% | 2% | -2% | -1% | 37% | -9% | -17% | -26% | -2% | -1% | -5% | -3% |
| G | -7% | 6% | -6% | -23% | 19% | -4% | 0% | 3% | -11% | 1% | 1% | 4% | 14% | 9% | -18% | 1% | -5% | -16% | -12% | 3% | -1% |
| H | 12% | -22% | -16% | -2% | 8% | 5% | 3% | 1% | -7% | 5% | -26% | -4% | 0% | -17% | -1% | 7% | -1% | -1% | 1% | -13% | -4% |

Agonist mode results from **Figure 2B**:

| | 1 | 2 | 3 | 4 | 5 | 6 | 7 | 8 | 9 | 10 | 11 | 12 | 13 | 14 | 15 | 16 | 17 | 18 | 19 | 20 | 21 |
|---|---|---|---|---|---|---|---|---|---|---|---|---|---|---|---|---|---|---|---|---|---|
| A | 2% | 4% | 0% | 5% | 0% | 1% | 0% | 0% | 3% | 1% | 1% | 0% | 0% | 1% | 0% | 1% | 1% | 1% | 1% | 0% | 2% |
| B | 5% | 2% | 0% | 0% | 0% | 2% | 1% | 1% | 9% | 4% | -4% | 5% | 7% | 1% | 3% | 1% | 0% | 6% | 0% | 3% | 1% |
| C | 2% | 0% | 3% | 5% | 3% | 0% | 4% | 1% | 0% | 2% | 1% | 12% | 1% | 4% | -1% | 1% | -3% | 0% | -2% | 2% | 0% |
| D | -2% | 1% | 0% | -2% | 0% | 3% | 3% | -4% | 1% | 4% | 1% | 5% | 1% | 0% | 1% | 2% | 3% | 5% | 2% | -1% | 1% |
| E | -3% | 4% | 1% | 4% | 1% | 44% | 14% | 0% | 0% | 0% | 0% | 1% | 7% | 4% | 3% | 4% | 0% | 8% | 4% | 0% | -1% |
| F | 1% | -1% | 1% | 2% | 1% | 1% | 1% | 3% | 0% | 3% | 1% | 0% | 13% | 1% | 1% | 2% | 0% | 1% | 1% | 1% | 1% |
| G | 2% | 1% | 5% | 0% | 0% | 0% | 9% | 0% | 0% | 3% | -11% | 0% | 7% | 0% | 3% | 0% | 0% | 9% | 3% | 2% | 3% |
| H | 1% | 2% | 3% | 2% | 0% | -2% | 3% | 1% | 3% | 1% | 4% | 0% | 5% | 4% | 2% | 1% | 2% | 0% | 0% | 0% | 1% |

**Table S3.** Impact of D2R point mutations on ML321 antagonist potencies as assessed using dopamine-stimulated Go BRET activation assays.

| Mutant D2R | Dopamine<br>Mutant $EC_{50}$/WT $EC_{50}$ ± SEM | ML321<br>Mutant $IC_{50}$/WT $IC_{50}$ ± SEM |
|---|---|---|
| $V91^{2.61}A$ | 1.23 ± 0.39 | **4.87 ± 1.2*** |
| $L94^{2.64}A$ | **3.34 ± 0.71*** | **2.24 ± 0.33*** |
| $W100^{EL1.50}A$ | **15.0 ± 5.8*** | **>40#** |
| $F110^{3.28}A$ | **24.7 ± 6.65*** | **0.20 ± 0.05*** |
| $T119^{3.37}A$ | ND | ND |
| $I184^{EL2.52}A$ | **36.8 ± 4.12*** | **1.77 ± 0.24*** |
| $S197^{5.46}A$ | ND | ND |
| $F390^{6.52}A$ | **9.1 ± 4.11*** | **>40#** |
| $H393^{6.55}A$ | **39.6 ± 13.9*** | 1.32 ± 0.26 |

Go-BRET activation assays were performed using the wild-type (WT) or mutant D2Rs as described in the Methods. Concentration-response curves (CRCs) were first performed for dopamine to determine the $EC_{50}$ and $EC_{80}$ values for each mutant receptor. Subsequently, cells expressing each mutant receptor were stimulated with an $EC_{80}$ concentration of dopamine and ML321 CRCs were performed to determine the $IC_{50}$ values for antagonism of this response. In each experiment, the dopamine $EC_{50}$ or ML321 $IC_{50}$ values for the mutant D2Rs were divided by their values for the WT D2R to assess the fold-change, if any, resulting from the mutation. The data shown represent the mean ± SEM values from 3-4 experiments each performed in triplicate. ND indicates that the mutation was severely deleterious to dopamine activity and thus not determinable. #Indicates that the ML321 CRC was incomplete at the highest concentration tested (10 μM) *$p < 0.05$ indicates significantly different from unity (1.0) as determined statistically using the one sample t test.

**Table S4.** Impact of D2R point mutations on ML321 binding affinities as assessed using [$^3$H]-methylspiperone competition binding assays.

| Mutant D2R | Spiperone Mutant Ki/WT Ki ± SEM | ML321 Mutant Ki/WT Ki ± SEM |
|---|---|---|
| V91$^{2.61}$A | ND | ND |
| L94$^{2.64}$A | 1.61 ± 0.23 | 1.96 ± 1.04 |
| W100$^{EL1.50}$A | ND | ND |
| F110$^{3.28}$A | 0.95 ± 0.23 | **0.05 ± 0.02*** |
| T119$^{3.37}$A | 2.04 ± 0.87 | 0.92 ± 0.31 |
| I184$^{EL2.52}$A | 0.99 ± 0.11 | 1.71 ± 0.34 |
| S197$^{5.46}$A | 0.86 ± 0.42 | 0.58 ± 0.1 |
| F390$^{6.52}$A | 1.36 ± 0.19 | **19.04 ± 5.2*** |
| H393$^{6.55}$A | 1.40 ± 0.38 | **9.77 ± 2.9*** |

Radioligand binding competition assays for spiperone and ML321 were performed using the wild-type (WT) or mutant D2Rs as described in the Methods. $IC_{50}$ values were determined for each competing ligand and used to calculate Ki values using the Cheng-Prusoff equation[50]. In each experiment, the ligand Ki values for the mutant D2Rs were divided by their values for the WT D2R to assess the fold-change, if any, resulting from the mutation. The data shown represent the mean ± SEM values from 3-4 experiments each performed in triplicate. ND indicates that the mutation was deleterious to [$^3$H]-methylspiperone binding and thus not determinable. *$p < 0.05$ indicates significantly different from unity (1.0) as determined statistically using the one sample t test.

**Supplemental Files:**

**ML321-D2R MD simulation.mpg**

**A representative molecular dynamics simulation trajectory showing the binding pose of ML321 in the D2R.** The carbon atoms of ML321 are in orange, while the receptor contact residues mutated in this study are in green. The sulfur atom of ML321's thiophene ring is in yellow. Note that the sidechains of I184$^{EL2.52}$, F389$^{6.51}$, and F390$^{6.52}$ (in spheres) tightly accommodate the tricyclic dibenzothiazepine moiety of ML321 in the orthosteric binding site, while the bulky phenyl sidechain of F110$^{3.28}$ pushes back and forth with the thiophene ring in the secondary binding pocket.
